# Dynamic changes in RNA-chromatin interactome promote endothelial dysfunction

**DOI:** 10.1101/712950

**Authors:** Riccardo Calandrelli, Lixia Xu, Yingjun Luo, Weixin Wu, Xiaochen Fan, Tri Nguyen, Chienju Chen, Kiran Sriram, Rama Natarajan, Zhen Bouman-Chen, Sheng Zhong

## Abstract

Chromatins are pervasively attached by RNAs. Here, we asked whether global RNA-chromatin contacts are altered in a given cell type in a disease context, and whether these alterations impact gene expression and cell function. In endothelial cells (ECs) treated by high-glucose and TNFα, we employed single-cell RNA-sequencing and *in situ* mapping of RNA-genome interaction (iMARGI) assay to delineate temporal changes in transcriptome and RNA-chromatin interactome. ECs displayed dramatic and heterogeneous changes in single cell transcriptome, accompanied by a dynamic and strong increase in inter-chromosomal RNA-DNA interactions, particularly among super enhancers (SEs). These SEs overlap with genes contributing to inflammatory response and endothelial mesenchymal transition (EndoMT), two key aspects of endothelial dysfunction. Perturbation of a high-glucose and TNFα-activated interaction involving SEs overlapping *LINC00607* and *SERPINE1* attenuated the pro-inflammatory and pro-EndoMT gene induction and EC dysfunction. Our findings highlight RNA-chromatin contacts as a crucial regulatory feature in biological and disease processes, exemplified by endothelial dysfunction, a major mediator of numerous diseases.

## Introduction

Mammalian genomes are extensively transcribed to RNAs, and a substantial portion of RNAs are physically associated with chromatins and thus are termed chromatin-associated RNA (caRNAs)^1^. Recent technological innovations enabled profiling of global RNA-chromatin interactions and revealed extensive RNA-chromosomal interactions in unperturbed cells^2–4^. Among these efforts, we have developed *in situ* mapping of RNA-genome interaction (iMARGI) assay which uses a bivalent linker to ligate to nuclear RNA and DNA followed by circularization for library construction^4, 5^. By utilizing the advantages of iMARGI, we observed that caRNAs are not only associated with the genomic sequences at or near their transcription sites, but are also attached to distal genomic sequences on the same or other chromosomes. A growing body of work focused on individual caRNAs suggests a modulatory role of these caRNAs in transcription through *cis* and *trans* regulation^6–8^. However, whether and how the global RNA-chromosomal contacts change in a given cell type that is undergoing a dynamic biological process, and whether they impact cell phenotype and function have not been explored.

Endothelial cells (ECs) lining the interface between circulating blood and vessel wall are crucial for the vital function of every tissue and organ through blood perfusion. Many pathological conditions such as hyperglycemia and chronic inflammation are detrimental to EC homeostasis^9–11^. The impaired EC function, generally defined as endothelial dysfunction, is typically manifested by a reduction of endothelial nitric oxide synthase (eNOS)-derived NO bioavailability and activation of pro-inflammatory responses^12, 13^. The chronic high glucose and pro-inflammatory cytokines [e.g. tumor necrosis factor alpha (TNFα)] can also induce endothelial to mesenchymal transition (EndoMT)^14, 15^, a process that provides a source of activated mesenchymal cells to contribute to endothelial dysfunction, pathological vascular remodeling, and a wide spectrum of cardiovascular diseases^16–21^. EndoMT is characterized by the suppression of EC markers (e.g. eNOS) and induction of mesenchymal markers [e.g. alpha smooth muscle actin (α-SMA)], with a distinct change in cell morphology^22, 23^. Although the importance of endothelial dysfunction and EndoMT have been well recognized in many diseases, the underlying molecular mechanisms remain incompletely understood. While it has become a consensus that chronic inflammation can promote EndoMT^13^, an outstanding question remains: how do the pro-inflammatory signals perpetuate to alter endothelial cell fate towards switching on mesenchymal gene program? Furthermore, how do hundreds of pro-inflammatory and pro-EndoMT genes encoded in distinct genomic loci across different chromosomes get activated in a concerted and coordinated fashion?

Super enhancers (SEs), the genomic regions comprising clusters of enhancers, have been attributed to drive transcription of genes involved in cell identity and cell-type specific functions^24, 25^. It is reasonable to postulate that a key aspect of endothelial dysfunction, i.e. EndoMT would engage activation of SEs that promote cell lineage conversion, which has not been tested. Moreover, many enhancers are actively transcribed into RNA transcripts, including long non-coding RNAs (lncRNAs)^26^. We have shown that these enhancer-derived lncRNAs can regulate endothelial gene transcription and function^8^. Thus we were interested in exploring the roles of SEs and SE-derived RNA, as well as their interactions in reprogramming the EC transcription underlying endothelial dysfunction and EndoMT. Testing this hypothesis by interrogating global RNA-chromatin interactome may reveal a novel regulatory feature contributing to endothelial dysfunction and EndoMT.

In this study, we first established a system where we induced EC dysfunction encompassing a robust pro-inflammatory activation and EndoMT phenotype. We then employed single-cell RNA sequencing (scRNA-seq) in conjunction with iMARGI to characterize the temporal changes in transcriptome with single cell resolution and RNA-chromatin interactome. We observed profound and dynamic activation of interchromosomal RNA-chromatin interaction especially among super enhancers (SE), the genomic regulatory regions crucial for cell function and identity. Many of the activated SEs that form interaction hubs encompass key regulators and mediators promoting inflammation, extracellular matrix (ECM) remodeling, and EndoMT. Among the highly activated SE networks, we identified an interaction involving SEs overlapping *LINC00607* (a novel long intergenic non-coding RNA with unknown function) and *SERPINE1*/PAI-1 (plasminogen activator inhibitor, a prominent regulator in endothelial dysfunction and many vascular diseases)^27^. Perturbing the RNA-chromatin contacts by *LINC00607* knockdown led to suppression of genes contributing to endothelial dysfunction and EndoMT, as well as attenuation of monocyte adhesion and EC senescence. Collectively, our study depicts a dynamic regulation of the RNA-chromatin interactome in endothelial dysfunction, a biological process closely implicated in numerous diseases.

## Results

### High glucose and TNFα induce a profound phenotypic change at cellular and molecular levels

We set out to establish and characterize an *in vitro* model to induce endothelial dysfunction. Given the prevalence of diabetes, one of the strongest risk factors for endothelial dysfunction preceding various diabetic complications^9^, we elected to subject human umbilical vein endothelial cells (HUVECs) to a prolonged treatment combining high glucose (HG, to mimic hyperglycemia) and TNFα (to mimic inflammation). Compared with ECs cultured under osmotic control (25 mM mannitol), the combined treatment (abbreviated as H+T) caused an evident switch from the classic endothelial cobblestone to mesenchymal-like spindle shape by 3 days, which became more distinguished by 7 days (Figure 1A). As a positive control for EndoMT, we also treated ECs with transforming growth factor beta (TGF-β) and interleukin 1 beta (IL-1β), which has been previously demonstrated to induce EndoMT within a similar time course^21, 28^. Both combinatory treatments provoked an apparent EC morphological change (Figure 1A), accompanied by the suppression of eNOS and induction of α-SMA (Figure 1B). Attendant with these transcriptional changes, α-SMA was also increased at the protein level by H+T as visualized by immunofluorescent staining, while CD31 remained to be expressed in ECs after the treatment (Figure 1C). Of note, H+T treatment induced a stronger EC morphological change and a higher induction of α-SMA (Figure 1A and 1B) and was used for the rest of this study.

**Figure 1.**
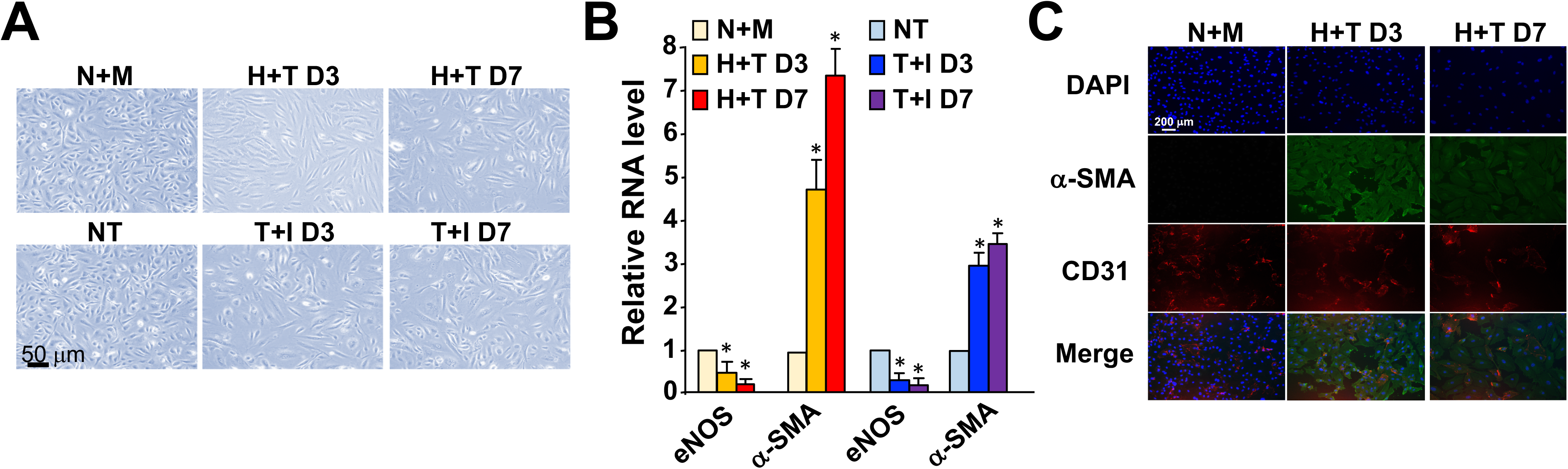
High glucose and TNFα induce a profound phenotypic change at cellular and molecular levels. HUVECs (at passage 5-6) were treated as follows: (top) by 25 mM mannitol as normal glucose and osmolarity control (N+M) or a combined treatment consisting 25 mM D-glucose and 5 ng/ml TNFα (H+T) for 3 or 7 days; (bottom) kept untreated (NT) or treated with TGF-β (10 ng/ml) and IL-1β (5 ng/ml) (T+I) for 3 or 7 days. (A) Cell morphology was monitored under bright-field microscopy. (B) mRNA levels of eNOS and α-SMA were quantified by qPCR with respective control set as 1. Data represent mean ± SEM from 3 independent experiments. * *P* < 0.05. (C) Immunofluorescent staining of α-SMA and CD31 and DAPI staining of the nuclei. Representative images from 5 independent experiments are shown in (A) and (C).

### Temporal changes in single EC transcriptome

Given the heterogeneity of ECs and the transitional nature of EndoMT^19, 22, 23^, we profiled the transcriptome in ECs subjected to the H+T treatment at the single-cell level using droplet-based single-cell RNA-sequencing (scRNA-seq). In ECs collected at three time points, i.e., Day 0 (osmolarity control; N+M), Day 3, and Day 7 (the latter two under H+T treatment), we identified more than 1,000 median genes per cell from over 4,000 cells sequenced per sample (Figures S1 and S2). Principal component analysis (PCA) of the highly variable gene expression revealed that ECs across three time points were not clustered separately, although a path following the time course was clear (Figure 2A). This implies that we have one EC population under different conditions, although there were clear variations in their transcriptomes (Figure 2A). When plotted in t-distributed stochastic neighbor embedding (t-SNE) plots, it is clear that ECs at the three different time points are identifiable as separate clusters (Figure 2B).

**Figure 2.**
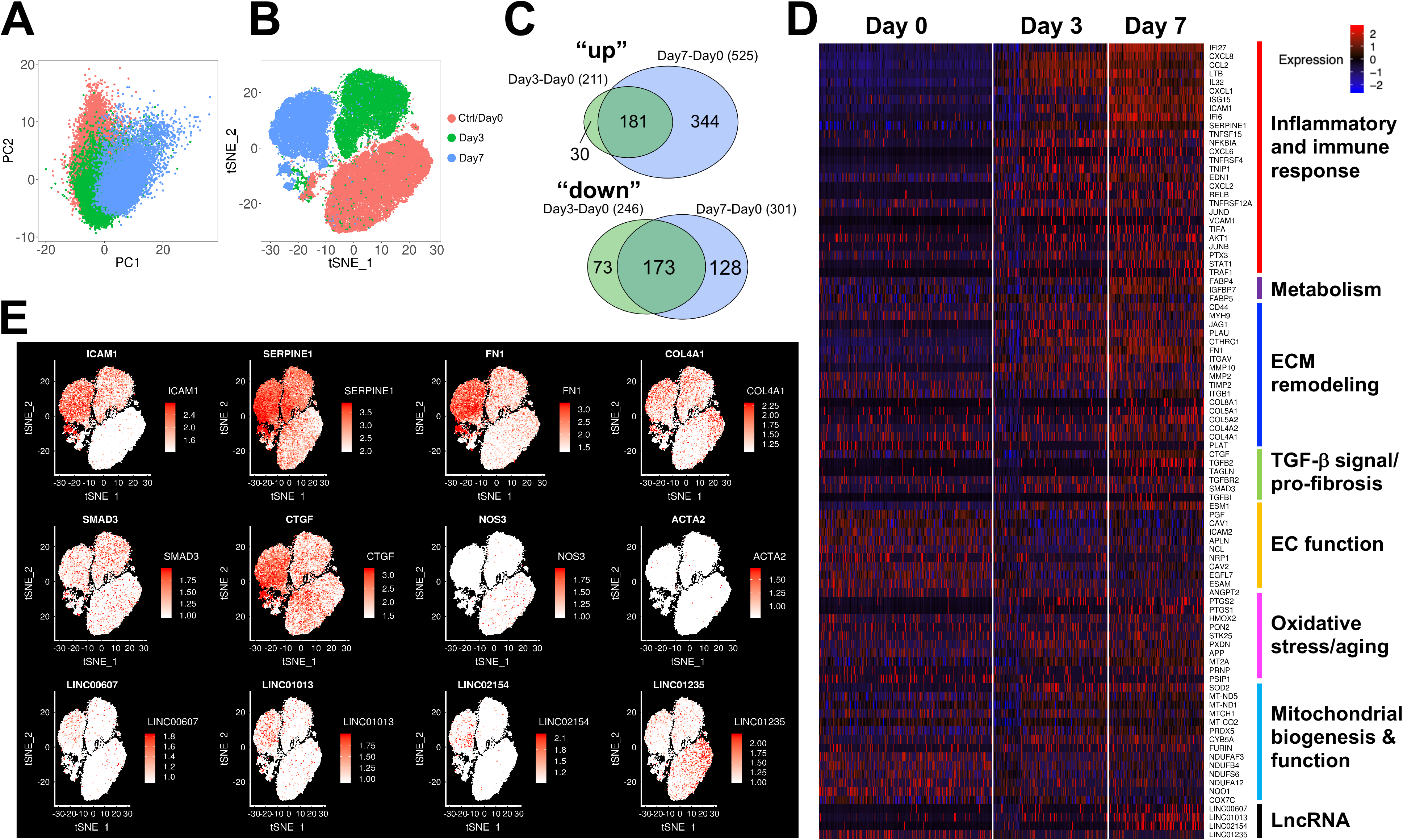
Temporal changes in EC transcriptome at single cell level. HUVECs in biological duplicates were treated as in Figure 1C and subjected to scRNA-seq. Populations with 4,000-15,000 cells were sequenced to generate >60M reads for each sample. (A) PCA analysis. Single cells are plotted in the first two principle components (PC) space and are label in red (Day 0, i.e. N+M), green (Day 3, i.e. 3-day H+T treatment) and blue (Day 7, i.e. 7-day H+T treatment). (B) t-SNE plot of the single cells with clear separation by condition into three distinct clusters. (C) Venn diagram of up- or down-regulated genes from scRNA-seq. Wilcoxon test was used, *P* < 9.72e-27 for Day 3 and *P* < 9.93e-115 for Day 7. (D) Heat map with RNA levels (represented by z-scaled values) of top DE genes in single ECs grouped into functional pathways. (E) t-SNE plots of the expression level of select genes in each single cell across the time course. The RNA levels are represented by log-normalized unique molecular identity (UMI) counts. See also Figures S1 and S2.

Next, we performed differential expression (DE) analysis on the single ECs among different time points. Using a threshold of 0.25 for the log2 fold change, we identified 457 genes differentially expressed between Day 0 vs Day 3 and 826 genes between Day 0 vs Day 7. Among these DE genes, 356 were common between Day 3 and Day 7, including 181 consistently upregulated and 173 consistently downregulated by H+T treatment (Figure 2C and S2).

To identify the biological processes and functional pathways involving these DE genes, we performed pathway enrichment analysis (see Table S1). We observed a significant enrichment of key pathways contributing to endothelial dysfunction and EndoMT, including inflammatory response [e.g. intercellular adhesion molecule (*ICAM1*), monocyte chemoattractant protein 1 (encoded by *CCL2*), chemokine ligand 2 and 8 (*CXCL2* and *CXCL8*), and plasminogen activator inhibitor 1 (PAI1, encoded by *SERPINE1*) were upregulated], ECM organization and remodeling [e.g. fibronectin (*FN1*) and collagens (*COL4, COL5, and COL8*) were upregulated], TGFβ signaling and fibrotic pathways [e.g. *TGFB1*, *TGFB2*, *SMAD3*, and connective tissue growth factor (*CTGF*) were induced], and oxidative stress response (Figure 2D). In addition to protein-coding genes, several long non-coding RNAs (lncRNAs) were also detected by scRNA-seq with DE patterns. These include the H+T-upregulated *LINC00607*, *LINC01013*, and *LINC02154*, and the downregulated *LINC01235*. Interestingly, *LINC00607* and *LINC01013* are respectively located upstream of *FN1* and *CTGF*, encoding two key regulators promoting fibrosis and EndoMT^15, 20^. The expression levels of these genes in the single ECs across the three time points are also depicted in Figure 2E.

Importantly, scRNA-seq data also revealed the heterogeneity and dynamics of EC transcriptomic changes. For example, the expression of EC hallmark gene eNOS (encoded by *NOS3*) was significantly decreased by H+T, evident not only by the reduced average mRNA level in single ECs, but also by the lowered percentage of ECs that express eNOS, following a time-dependent manner. In the same time course, pro-inflammatory and pro-fibrotic genes (e.g. *ICAM1*, *FN1*, and *SERPINE1*) were increased in mRNA levels in single ECs and in percentage of ECs with positive expression across time (Figure 2D and 2E). We also observed varied patterns of transcriptional changes among DE genes engaged in different molecular pathways and cellular functions. The most robust induction was seen in the pro-inflammatory genes. For instance, *ICAM1* was detected in <2% of control ECs at Day 0, while it was expressed in 68% of ECs at Day 3 and up to 89% of ECs at Day 7. Similar but less drastic dynamics were observed for genes involved in ECM organization and remodeling. For example, *FN1* was expressed in 36% of control ECs, which increased to 73% of cells by Day 3 and then further to 95% of cells by Day 7. In contrast, the mesenchymal marker α-SMA was induced in a much slower pattern, i.e. from 0.2% of control ECs to 1.4% of cells after 7 days of H+T treatment (Figure 2E). These results suggest a time-dependent signaling cascade initiated by a strong inflammatory response, which relays to substantial ECM remodeling and eventually perpetuates EndoMT.

### Dynamic changes in global RNA-chromatin interactome in EC dysfunction

With the characterized transcriptome of ECs undergoing the H+T treatment, we proceeded to profile global RNA-chromatin interactions in this dynamic process using iMARGI. To test the possibility that H+T may induce global chromatin conformation changes in ECs to change RNA-chromatin contacts, we first performed Hi-C to map the chromatin architecture in ECs under the prolonged treatment to induce endothelial dysfunction. Based on the Hi-C data, there was no significant change among either the intra- or inter-chromosomal interactions (Figure 3, Table S2).

**Figure 3.**
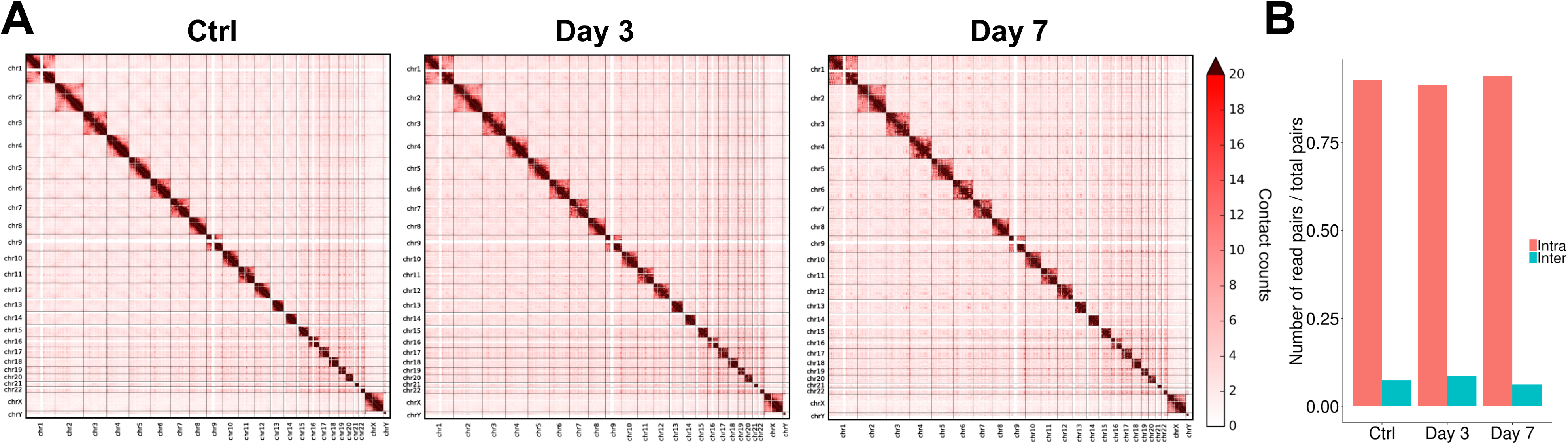
Lack of changes in EC chromatin conformation revealed by Hi-C. (A) Pairwise heat maps of Hi-C contacts plotted with normalized contact counts. (B) Bar plot showing the percentage of intra- and inter-chromosomal read pairs in three conditions of ECs.

In contrast, iMARGI results from the same EC populations showed dramatic changes in RNA-chromosomal interactions. Among over 65 million total read pairs, we detected substantial changes in the intra- as well as inter-chromosomal interactions. Furthermore, the intra-chromosomal RNA-DNA interactions were decreased, whereas the inter-chromosomal interactions were remarkably increased in a time-dependent manner by H+T treatment (Figure 4A and 4B; Table S3).

**Figure 4.**
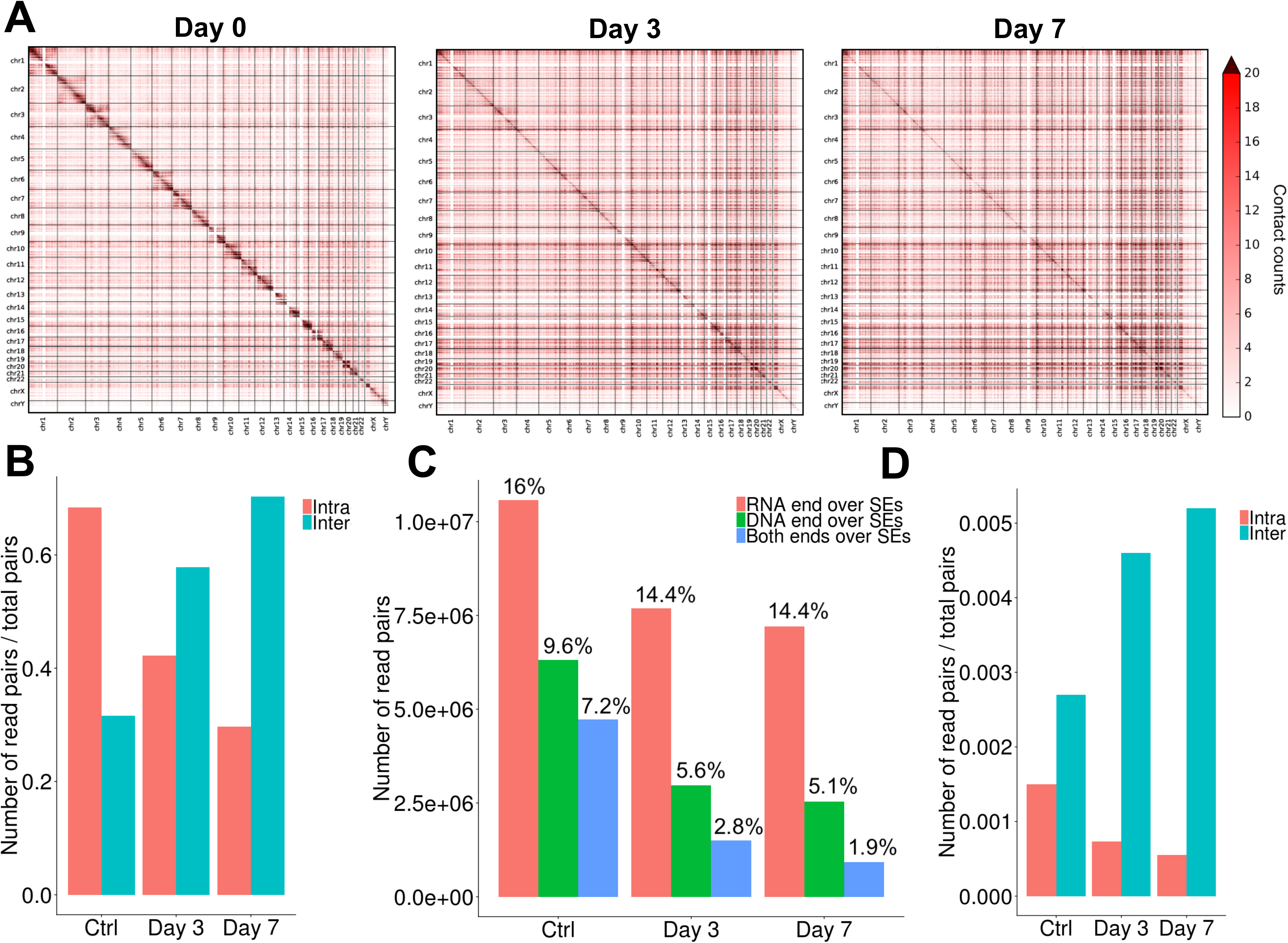
iMARGI revealed dynamic changes in RNA-DNA interactions in ECs. (A) Global iMARGI heat maps at 1 Mb resolution. Contact counts were scaled by the total reads in each sample. (B) Bar plot of the number of intra- and inter-chromosomal read pairs scaled by the total reads in ECs from three time points. (C) Bar plot of the SE coverage in the MARGI data. The percentages above each bar are in respect to the total number of read pairs in each sample (i.e., control/Day 0, Day 3 and Day 7). Note the enrichment of read pairs in SEs considering the occupancy of SEs in the entire genome, i.e. 3.1%. (D) Bar plot of the number of intra- and inter-chromosomal read pairs derived from super enhancers (SEs) relative to the total reads in ECs from three time points.

### Activation of inter-chromosomal interactions among super enhancers in EC dysfunction

Because we observed more DE genes upregulated than downregulated by Day 7 of H+T treatment (Figure 2C), we argued that the increased inter-chromosomal RNA-chromatin interactions, rather than the decreased intra-chromosomal interactions, would be more directly related to the transcriptional induction of genes promoting EC dysfunction. We next asked whether the increased inter-chromosomal RNA-DNA interactions were enriched in certain genomic regions. Given that SEs control gene transcription in cell-type specific processes and transient TNFα has been shown to induce reorganization of SEs to activate endothelial inflammation^24, 25, 29^, we assessed the degree of RNA-DNA interactions derived from SE regions. To do so, we leveraged the previously annotated SEs in HUVECs (dbSUPER)^30^. These SEs span around 94.5 million bp in total, which is equivalent to approximately 3% of the entire human genome. Among the total read pairs revealed by iMARGI, 20-30% involve SEs (either with the RNA or DNA end, or both ends stemming from SEs) (Figure 4C), suggesting an enrichment of RNA-chromosomal interactions in SE regions.

Similar to the patterns observed for the whole genome (Figure 4A and 4B), the RNA-DNA interactions involving SEs also showed a decrease intra-chromosomally and an increase inter-chromosomally as the H+T treatment prolonged (Figure 4D, 5A and 5B, and Figure S3). In order to analyze this phenomenon, we strictly selected only highly interacting SEs. This was achieved by setting a threshold *gamma* on the number of read pairs among SEs scaled by the total pairs in each sample to avoid any bias due to the different sample sizes. At a very restrictive *gamma* of 2e-7 (Figure S4), the inter-chromosomal interacting SE pairs increased from 510, to 2234, and further to 3367, reaching a 6.6-fold change by Day 7 as compared with control (Figure 5B). Notably, the tendency of increasing number of SE pairs was independent from the choice of the threshold, and it became even stronger as we increased the threshold (Table S4). Conversely, with the same *gamma* of 2e-7, the intra-chromosomal interacting SE pairs decreased from 1,609 in control to 789 in Day 3 and 621 in Day 7 (i.e., 2.6-fold decrease as compared with control). Moreover, this tendency remained relatively steady and became weaker at higher thresholds (Table S4). This observation, together with the fact that SEs are by nature associated with gene activation, prompted our focus on the increase in SE-derived inter-chromosomal interactions for the rest of this study.

**Figure 5.**
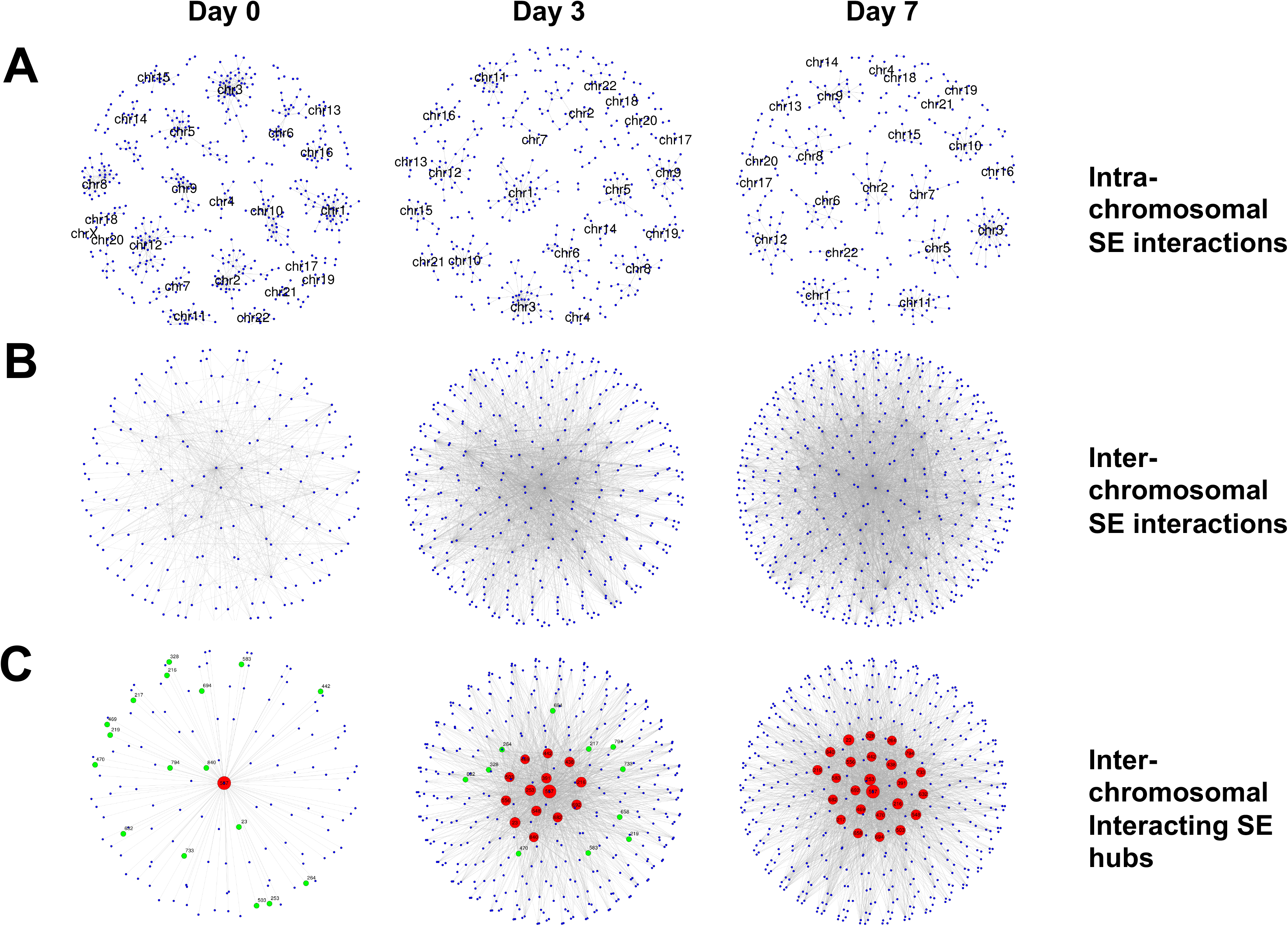
H+T increases inter-chromosomal interactions among SEs. (A) Intra-chromosomal and (B) Inter-chromosomal RNA-DNA interactions among SEs represented as networks (*gamma* = 2e-7). Each node is a SE and edges connect interacting SEs. (C) Inter-chromosomal networks with SE hubs with degree of nodes (DON) > 60 are highlighted in red. Each hub is labeled with a SE index (see Table S5) and has a dimension proportional to its DON. The 14 hubs in Day 3 are a subset of the 25 hubs in Day 7. Green nodes are normal nodes in control (Day 0) and Day 3 samples but became SE hubs by Day 7. See also Figures S3-S5.

In the subsequent effort, we queried whether our inter-chromosomal networks comprise SEs with a significant high number of interactions (edges), which may play a more meaningful role in the cellular transition (i.e. inflammation to EndoMT) process. To this end, we strictly defined SE hubs as those with a degree of node (DON) above 60 (over the 95^th^ percentile values of the DON distributions for the three conditions) (Figure S5). Given this definition, there was only 1 SE hub (which overlaps with *MALAT1*) identified in control (Day 0) ECs, with 131 active SE nodes and 130 interacting SE pairs. This is in line with previous reports from us and others, wherein *MALAT1* is engaged in broad trans-chromosomal interactions in several unperturbed cell types^2, 4, 31, 32^. After 3 days of treatment, the number of hubs had increased to 14, with 382 active SE nodes and 1,738 interacting SE pairs. After 7 days of treatment, while all 14 active SE hubs remain as a subset of all the SE hubs, 11 novel hubs emerged, thus giving rise to 25 active hubs in total, with 456 participating SE nodes and 2,627 interacting SE pairs (Figure 5C).

Collectively, H+T treatment induced a dramatic increase in inter-chromosomal RNA-DNA interactions particularly among SEs, with a set of SE hubs forming novel and strong interactions among each other in ECs.

### Induction of novel *SERPINE1-LINC00607* SE interactions in ECs by high glucose and TNFα

We reasoned that the interactions involving the emerging SE hubs would be most relevant to H+T-induced changes, including pro-inflammatory activation, ECM remodeling, and EndoMT. Strikingly, among the 15 SE hubs that emerged by Day 3 and remained active by Day 7, SE hub #391 overlaps with *SERPINE1* (Figure 6A, Table S5), a key regulator linking hyperglycemia, inflammation, and EndoMT^27, 28^ and one of the top H+T-induced genes in ECs identified by scRNA-seq (Figure 2D and 2E). Furthermore, SE #145 overlapping with *LINC00607* (one of the lncRNAs significantly induced by H+T) (Figure 2D and 2E), became an active SE node by Day 3 and formed a novel interaction with the *SERPINE1* SE by Day 7 (Figure 6A). This is evident by the increase in reciprocal interactions between LINC00607 RNA-*SERPINE1* DNA and SERPINE1 RNA-*LINC00607* DNA (Figure 6B-D).

**Figure 6.**
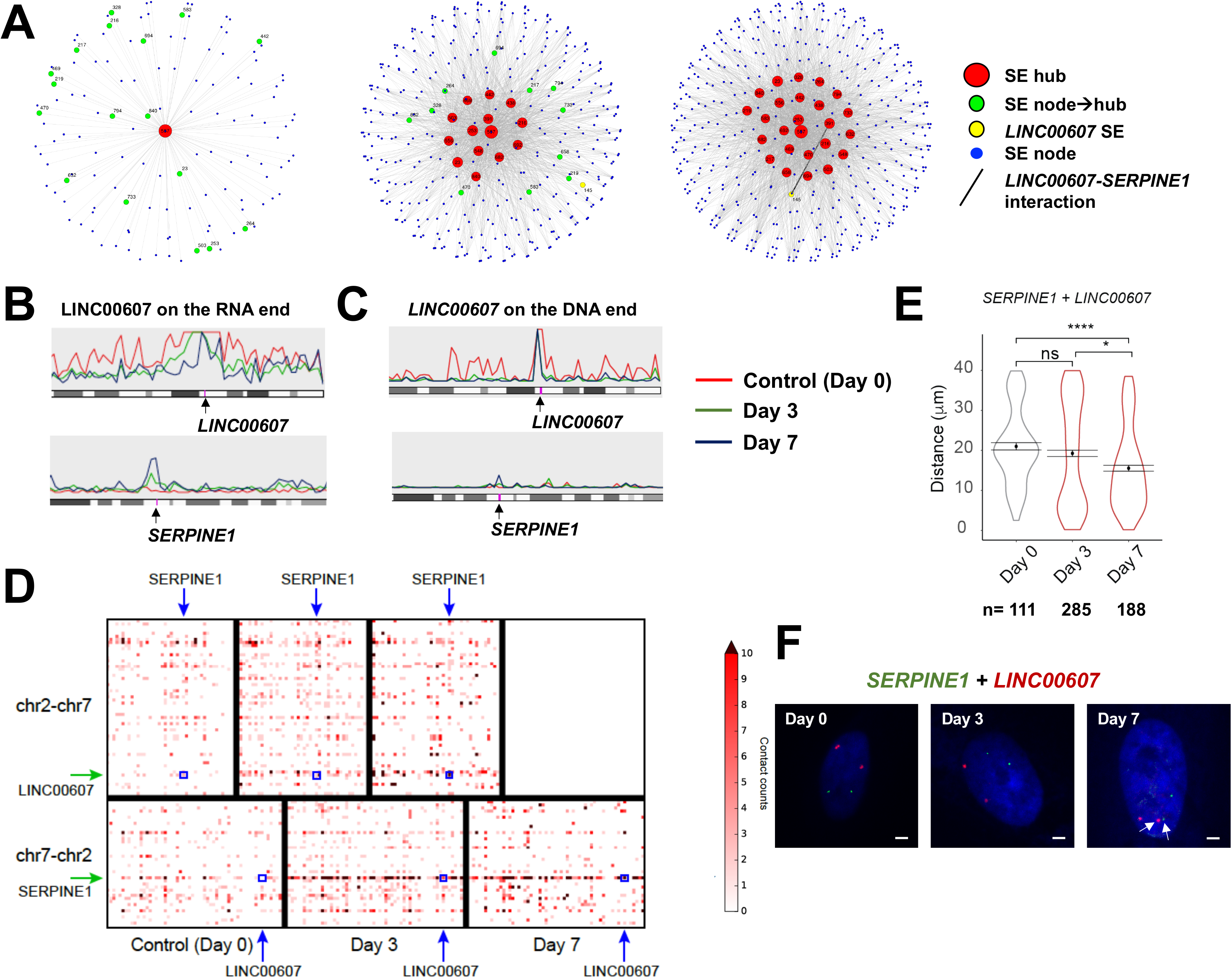
*SERPINE1* and *LINC00607* form novel SE interaction by H+T treatment. (A) SE hub interaction network across three conditions. *LINC00607* SE is plotted bigger and in yellow. Note the presence of interactions between *LINC00607* (SE index #145) and *SERPINE1* (SE index #391) only in Day 7 ECs (indicated by the bold edge) but not in control (*LINC00607* SE not present as a node) and Day 3 (*LINC00607* SE showing up as a node but not interacting with SERPINE1). (B, C) Snapshots from coverage plot of the iMARGI-revealed LINC00607 RNA targets (in B) and RNAs targeting the *LINC00607* locus (in C). Note the increased interactions between LINC00607 RNA and *SERPINE1* DNA (B) and *LINC00607* DNA and SERPINE 1 RNA (C) across time course. (D) Heat map showing LINC00607-SERPINE1 reciprocal interactions. Each pixel is a SE. Top: LINC00607 on the RNA end and *SERPINE1* on the DNA end; Bottom: *LINC00607* on the DNA end and SERPINE on the RNA end. (E) Distance between *SERPINE1* and *LINC00607* enhancer regions measured by DNA FISH. Total number of pairs are listed on the bottom of the plot. p-values generated by nonparametric Wilcox-test with Bonferroni correction for multiple comparisons. **** *P* < 0.0001. ns = not significant. (F) Representative images of DNA FISH. Nuclei were stained by DAPI; bar= 2 μm. Arrow heads indicate example proximal pair. See also Figure S6 and S7.

The increase in RNA-DNA interactions involving *SERPINE1* and *LINC00607* SEs prompted us to test whether these two chromosomal loci became closer upon H+T treatment. DNA FISH probes were designed against the interaction-enriched chromosomal regions spanning *LINC00607* and *SERPINE1* enhancers. Indeed, the mean distance between *LINC00607* and *SERPINE1* loci was slightly shortened by Day 3, and significantly reduced by Day 7 of H+T treatment (Figure 6E and 6F). In contrast, DNA FISH signals from *SERPINE1* and the non-interacting control region on chromosome 2 did not show any significant changes in their distance (Figure S7).

### Perturbing LINC00607 network inhibits transcriptional activation of genes promoting EC dysfunction

Next, we asked whether the activation of SE-derived inter-chromosomal interactions has any regulatory role in EC dysfunction. To answer this question, we intended to perturb the *LINC00607-SERPINE1* axis through disrupting the RNA-chromosome interaction. We reasoned that perturbing SE overlapping with SERPINE1 will result in effects that are most likely due to the perturbation of SERPINE1, rather than the RNA-chromosomal interaction. Thus, we opted to inhibit the lncRNA transcripts derived from *LINC00607* SE, which would disrupt the RNA-DNA interactions between *LINC00607* and *SERPINE1* SEs. To do so, we tested two locked nucleic acid (LNA) Gapmers targeting different regions of LINC00607 RNA (Figure 7A). Although both LNA Gapmers efficiently decreased the level of LINC00607 in total EC extracts and cytoplasmic fraction, only LNA1, but not LNA2, led to a decreased level of LINC00607 RNA in the nucleus (Figure 7B).

**Figure 7.**
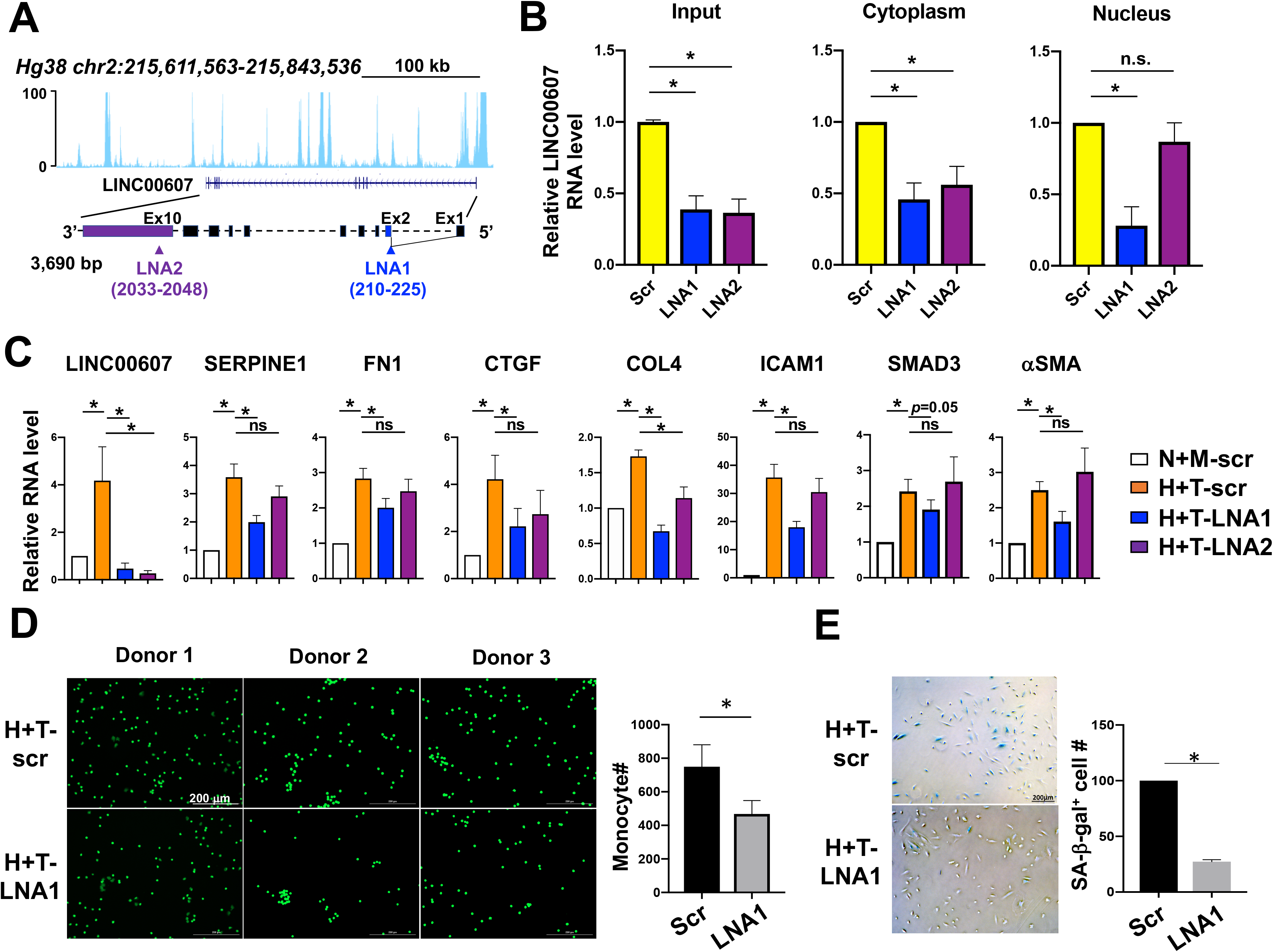
Inhibition of LINC00607 attenuates molecular and cellular changes contributing to endothelial dysfunction. (A) Illustration of *LINC00607* genomic locus and gene structure and LNA Gapmers targeting of LINC00607 RNA. (B) qPCR detection of LINC00607 in EC subcellular fractions. (C) HUVECs were transfected with LNA with scramble (scr) or LINC00607 targeting sequences before H+T treatment or kept under control condition. qPCR analysis of RNA levels of indicated genes. (D, E) ECs transfected with scramble or LNA1 and then treated by H+T were used in (D) monocyte adhesion assay performed with fluorescently labeled peripheral blood-derived monocytes from individual donors and in (E) SA-β-gal assay. The number of attached monocytes in (D) and ECs with positive β-gal staining in (E) were quantified. Data represent mean ± SEM from 4-6 independent experiments.

When we used these LNAs to inhibit LINC00607 in ECs treated by H+T, only LNA1 significantly suppressed the induction of SERPINE1. Additionally, we examined other markers related to inflammation, ECM remodeling, and EndoMT as revealed by scRNA-seq. Interestingly, LNA1 also suppressed the induction of FN1, COL4, ICAM1, SMAD3 and α-SMA (Figure 7C). In contrast, LNA2 resulted in no significant effect in these markers, with the exception of COL4 (Figure 7C), suggesting the effect of LINC00607 on the transcription of these genes preferentially occurred in the nucleus, likely through the chromatin-associated RNA transcripts.

To examine the functional consequence of perturbing LINC00607-SERPINE1 axis in ECs, we performed monocyte adhesion assay to assess the inflammatory state of ECs. The inhibitory effect of LINC00607 LNA1 in monocyte adhesion to ECs was consistent using peripheral blood-derived monocytes from 4 different donors (Figure 7D). Additionally, we performed the senescence-associated β-galactosidase (SA-β-gal) assay to evaluate EC senescence^33^, an aspect of EC dysfunction^34^. In line with the gene expression changes and monocyte adhesion assay, SA-β-gal staining of ECs was reduced by LINC00607 LNA (Figure 7E), collectively supporting the role of LINC00607 and its engaged SE network in promoting endothelial dysfunction.

## Discussion

The organization of 3D genome is an area under intensive investigation. While existing studies provide perspectives largely on DNA-DNA interactions in gene regulation, there is much less known about how RNA-chromatin contacts contribute to functional regulation in a given biological process. In the context of endothelial dysfunction, we characterized the dynamic changes in previously unexplored RNA-chromatin interactome in relation to transcriptome. Our major findings include: 1) the combined H+T treatment induced a profound and dynamic remodeling in global RNA-chromatin interactome enriched among SEs, whereas the chromosomal conformation remained fairly constant; 2) there was a time-dependent increase in inter-chromosomal interactions in ECs undergoing dramatic transcriptional changes, evident by the robust increase in the numbers of SE hubs and the degree of interactions with other SEs; 3) the activation of SE hubs and networks contributes to the induction of genes promoting inflammation, ECM remodeling, and EndoMT, and perturbation of the identified *LINC00607*-*SERPINE1* SE interaction suppressed the expression of these genes that contribute to EC dysfunction. Collectively, our study presents an integrative approach that provides novel insights into how RNA-chromatin contacts may alter and function as a critical regulatory layer in gene expression and vital disease-relevant processes.

The robust increase in inter-chromosomal interaction enriched among SEs was unexpected and intriguing. Although intrachromosomal interactions are often dominant in the chromatin architecture, inter- or trans-chromosomal interactions can also occur, even at comparable frequency and may result in functional regulation^7, 8, 35^. Monahan et al recently reported that olfactory receptor gene clusters make specific and robust inter-chromosomal contacts orchestrated by intergenic enhancers. These contacts increase with cell differentiation and transcriptional activation of olfactory receptor gene expression, providing functional support for the role of trans-chromosomal interactions^36^. In light of these findings, our results suggest that the chromatin remodeling involving trans- or inter-chromosomal interactions may synergize with RNA-chromosomal interactions. Among the SE hubs activated by H+T (Table S5), many overlap with genes that have been shown to be induced at the transcriptional level and play pivotal roles in endothelial dysfunction and EndoMT. In addition to *SERPINE 1*, runt-related transcription factor 1 (*RUNX1)* is a key transcription factor (TF) induced by high glucose, and its inhibition has been shown to protect against adverse cardiac remodeling^37, 38^. SMAD3 is a key TF driving EndoMT^16^, von Willebrand factor *(VWF*) has long been recognized as a marker for endothelial dysfunction^39, 40^, thrombospondin (*THBS1*) is a multifunctional extracellular matrix molecule that promotes inflammation and apoptosis^41^, and *PVT1* has been shown to contribute to the susceptibility of diabetic nephropathy, in which EC dysfunction is essential^42^. The dynamic and sustained activation of these SE hubs and the associated SE networks suggests a novel molecular mechanism at the RNA-chromatin contact level underlying their reported central role in EC dysfunction and vascular diseases.

Notably, many of the SE that became hubs by Day 3 or Day 7 were already active in control ECs. For example, SE #391 (overlapping SERPINE1) was an active node in control ECs, and became a SE hub by Day 3 and further nucleated a higher dimension of interacting network by Day 7. However, SE #145 (overlapping *LINC00607*) did not appear as an active node in the SE network in control ECs, but became active by Day 3 and formed a novel interaction with *SERPINE1* SE. Within the same time frame, the transcription of LINC00607 RNA increased from <3% of control ECs to over 21% of treated-ECs. A similar pattern was observed for SE #692, which overlaps with SMAD3 (Figure 5C and Table S5). Based on these dynamics, it is reasonable to speculate that, within a given cell type, while a subset of SEs is key to the maintenance of cell identity, additional SEs may become activated by external stimuli to form new hubs for RNA-chromatin interactions. In the scenario of EC dysfunction, the activation of these stimuli-dependent SE hubs may activate the transcription of genes switching on inflammation and ECM remodeling pathways (e.g. SERPINE1), and, with the growing SE hubs and expansion of SE networks, amplify the transcriptional activation of genes promoting EndoMT (e.g. *SMAD3* and *LINC00607*). While our findings are in line with the previous study demonstrating strong activation of inflammatory SEs in ECs transiently treated with TNFα (for 1 hr), the strong and sustained activation of SE hubs overlapping *SERPINE1* (key regulator in ECM remodeling) and SMAD3 (key transcription factor for EndoMT) provides insights into how the initial activation of inflammation promotes ECM remodeling and EndoMT in a prolonged time course. These changes in RNA-chromosomal contacts provide temporal and spatial explanation regarding how these molecular and cellular events are coordinated in the nucleus.

How does high glucose and TNFα induce such a remarkable increase in inter-chromosomal interactions? Examining the size of nuclei, we did not observe any significant difference due to the treatment (Figure 6F). The lack of changes in Hi-C maps suggests that it is unlikely that the changes in global chromatin architecture drove the remodeling of RNA-chromatin interactome. We also tested whether this process involves phase transition by staining for SC35, a marker for nuclear speckles, which revealed a lack of demonstrable changes in the SC35 pattern in ECs (Figure S8). In contrast, we detected shortened distance between the select SEs, exemplified by SEs overlapping *LINC00607* and *SERPINE1* in ECs at Day 7 (Figure 6F), implicating that RNA-chromosomal interactions may form due to the activation of SEs, which may in turn promote inter-chromosomal DNA-DNA interaction in a cell-type- and stimulus-specific manner.

Using *LINC00607-SERPINE1* SE axis as an example, we demonstrated that nuclear inhibition of LINC00607, conceptually by perturbing the RNA-chromosomal interactions between the *LINC00607* and *SERPINE1* SEs resulted in significant suppression of SERPINE1, of which the DNA region forms direct proximity contacts with LINC00607 RNA (and vice versa) (Figures 6 and 7C). Such inhibition also decreased the expression of *FN1* encoded downstream of LINC00607 SE, which is likely due to a *cis*-regulation. Additionally, several other genes promoting EC dysfunction and EndoMT, e.g. *SMAD3, COL4*, and *CTGF* were also suppressed. Using a two-step network approach to explore secondary effects and multi-way SE interactions based on the iMARGI data, we found that, although LINC00607 did not show any interaction in the control ECs, it formed a network of inter-chromosomal contacts by Day 3 (with 779 interacting SE pairs consisting of 375 SE nodes), whose dimension further increased by Day 7 (with 1,863 interacting SE pairs consisting of 443 SE nodes) (Figure S9 and Table S6). Of note, SEs overlapping with *SMAD3*, *COL4*, and *CTGF* were present in the *LINC00607* SE-derived network, suggesting that chromatin-associated LINC00607 RNA may be an essential component of the multi-SE-orchestrated transcriptional hubs.

Finally, the dynamics in RNA-chromatin interactomes in endothelial dysfunction may have implications in our understanding of diseases and development of therapeutic strategies. For example, high glucose may induce a “metabolic memory” (i.e. the changes persist after the switch of hyperglycemia to normoglycemia which may underlie the persistent damage due to diabetic conditions (e.g. hyperglycemia and chronic inflammation)^43–45^. Interestingly, replenishing ECs with control media after 3 days of H+T treatment partially reversed the cell morphology change (Figure S10A). However, at the molecular level, whereas the H+T suppressed eNOS and ICAM1 mRNA levels could be almost completely reversed, those of LINC00607, SERPINE1, FN1, and CTGF could not be reversed. In fact, the induction of LINC00607, SERPINE1, and FN1 continued or sustained despite the removal of detrimental stimuli (Figure S10B). These data suggest that the mechanisms involving the RNA-chromatin contacts may be an important, yet poorly elucidated, layer underlying various biological processes, as has been proposed recently^46, 47^. As proof-of-principle, our study provides a first example of how mapping of RNA-chromatin interactomes can provide novel insights into understanding endothelial dysfunction, a dynamic and vital process preceding a variety of diseases.

## Material and Methods

### Cell lines

HUVECs (Passages 5-8) from pooled donors were used in this study. The cells have been tested negative for mycoplasma contamination and prescreened to demonstrate stimulation-dependent angiogenesis and key EC signaling pathways. ECs were cultured at 37°C with 5% CO_2_ in M199 (Sigma M2520) supplemented with 20% FBS (Hyclone, SH30910.02), β-endothelial cell growth factor (Sigma, E1388), and 100 units/ml penicillin and 100 mg/ml streptomycin (Thermo Fisher Scientific).

### Cell transfection and stimuli

Two antisense LNA GapmeRs specifically targeting two different regions of LINC00607 (NR_037195.1) were designed and purchased from QIAGEN (see table below for sequences). LNAs were separately transfected into HUVECs with Lipofectamine RNAiMAX following the protocol provided by the manufacturer. The cells were cultured in M199 complete medium 4-6 hr after transfection, and then subjected to high glucose and TNFα treatment. High glucose condition was generated by adding D-glucose into the culture media to a final concentration of 25 mM. TNFα was added to the culture media to a final concentration of 5 ng/mL. 25 mM D-mannitol was used as an osmolarity control. Typical EndoMT stimuli TGF-β and IL-1β were added to the culture media at a final concentration of 10 ng/mL and 1 ng/mL, respectively.

### RNA extraction and quantitative PCR (qPCR)

Total RNA was isolated using TRIzol reagent. cDNAs were synthesized using PrimeScript™ RT Master Mix containing both Oligo-dT primer and random hexamer primers. qPCR was performed with Bio-Rad SYBR Green Supermix following the manufacturer’s suggested protocol using the Bio-Rad CFX Connect Real Time system. All primer sequences used in qPCR amplification are listed in Table S7.

### Immunofluorescent staining

HUVEC cells were plated on coverslips (pre-coated with poly-L-lysine and 0.1 M collagen solution), let adhere, and treated with high glucose and TNF-α or cultured in control conditions. For α-SMA and CD31 immunostaining, cells were washed with PBS and fixed with 4% formaldehyde diluted in PBS for 15 min at room temperature (RT). The fixed cells were rinsed three times by PBS for 5 min each, and incubated in a blocking buffer containing 5% BSA in PBST (PBS with 0.1% Triton X 100) for 1 hr. A phycoerythrin (PE)-conjugated rat anti-CD31 antibody (at 1:500 dilution) or a rabbit anti-human α-SMA were added to cells in blocking buffer at 1:500 and 1:1000 dilution and incubated overnight at 4°C. From this step on, cells were protected from light. After rinsing in PBST for three times (10 min each), cells were incubated with a FITC-conjugated goat anti-rabbit antibody (for α-SMA staining) at 1:750 dilution in blocking buffer at RT for 1 hour. After three times washes in PBST for 10 min each, cells were stained with DAPI at 1:3000 dilution in PBST at RT for 15 min. The fluorescence images were taken with an Echo revolve reverse fluorescence microscope.

For SC35, cells were fixed with 4% paraformaldehyde in PBS (pH = 7.2) (RT, 30 min) and then rinsed with PBS. Cells were permeabilized with 0.1% Triton-X100 in PBS (PBST), 15 min, RT. The cells were blocked with 5% BSA in PBST for 30 min at RT and incubated with primary antibody (monoclonal mouse anti human SC35, Abcam) at 1:250 dilution for 1 hr at 37°C. Subsequently, cells were incubated with secondary antibody (Alexa Fluor 568-conjugated goat anti mouse IgG, Invitrogen) at 1:500 dilution in blocking buffer at 37°C for 30 min, followed by a final mounting with 6 μL ProLong Glass Antifade mountant with NucBlue Stain. Images in size of 512 × 512 were acquired on Applied Precision OMX Super Resolution Microscope using a 100X/1.518 oil objective (GE Healthcare Life Sciences) (pixel size = 0.079 μm). A series of z-stack images across the cells were acquired with the thickness of 0.3 μm per section.

### Single-cell RNA-seq and data analysis

Mannitol control (Day 0), HG+ TNF (Day 3), and HG+ TNF (Day 7) with duplicates were prepared as single cell samples for sequencing using Drop-seq protocol with 10x Genomics Chromium System at the Integrative Genomics Core at City of Hope. There are >60M reads/sample, 4,000-15,000 cells/sample. scRNA-seq data have been processed using the standardized pipeline provided by 10X Genomics (v3.0) and aligned to human hg38 reference transcriptome. The R package Seurat (v2.3.4) was used to analyze scRNA-seq data following published guidelines^48^. First, we performed a filtering step using well established quality control metrics. Rare cells with very high numbers of genes (potentially multiplets) as well as high mitochondrial percentages (low quality or dying cells often present mitochondrial contamination) were removed. We set the upper-threshold for both of those features as the 99th percentile of their distribution in each sample (Figure S1). In addition, cells exhibiting a gene count lower than 300 were filtered out as potential low-quality cells or empty droplets. Filtration led to the removal of ~2% of the total cells (from 60,841 to 59,605 total cells across all the samples).

Next, data were normalized by default in Seurat. We employed the global-scaling normalization method “LogNormalize” that normalizes the gene expression measurements for each cell by the cell total expression, multiplies it by a scale factor (10,000 by default), and log-transforms as log(x+1) the result. We selected highly variable genes (HVGs) and scaled the gene expression data for downstream analysis. HVGs were calculated as default in Seurat by using the log(variance to mean ratio) (logVMR) for each gene in the dataset. Genes were then sorted by decreasing logVMR, and then we extracted the top 1,000 HVG. Unwanted sources of variation, such as mitochondrial expression and number of detected molecules per cell, were regressed out and the expression of each HVG was scaled to obtain a z-score for each gene across all the single cells in the dataset. Principal component analysis (PCA) was performed across all cells and the top 1,000 HVGs using the scaled z-scored expression values. The first 20 significant PCs were then used as input to the t-SNE algorithm.

Differentially expressed (DE) genes analysis was performed using the non-parametric Wilcoxon test (default in Seurat and one of those that globally perform the best according to Soneson et al)^49^. The test was performed using default parameters in Seurat. Thus, only genes expressed in at least 10% of the cells in a sample were used in the DE genes analysis. To extract DE genes, the threshold for the log fold-change of the average expression (logFC_avg) between two samples was left to 0.25 (either for up- or down-regulated genes). We used a pseudocount of 1 (as default) to be added to the averaged expression values when calculating logFC. This prevents extremely-lowly-expressed genes from dominating the differential expression analysis.

### Hi-C

Hi-C was performed using an Arima-HiC kit (Arima Genomics, Inc.) following the manufacturer’s manual. Hi-C data were processed and plotted using HiCtool v2.1^50^. Data were normalized using the matrix balancing approach performed by Hi-Corrector ^51^ and incorporated into HiCtool.

### *In situ* MARGI (iMARGI) assay and data analysis

*In situ* MARGI was performed as described in our recent report^5^. Briefly, iMARGI started with crosslinking cells using 1% formaldehyde, collecting nuclei, followed by fragmenting RNA and DNA in nuclei using RNase I and restriction enzyme AluI. A specifically designed linker sequence was introduced to the permeated nuclei to ligate with the fragmented RNA and subsequently ligate with spatially proximal DNA. After the ligation steps, nuclei were lysed and crosslinks were reversed. Nucleic acids were purified and subsequently treated with exonucleases to remove any linker sequences that were not successfully ligated with both RNA and DNA. The desired ligation products in the form of RNA-linker-DNA were pulled down with streptavidin beads. The RNA part of the pulled down sequence was reverse transcribed into cDNA, resulting in a complementary strand of (5’) DNA-linker-cDNA (3’). Single-stranded DNA-linker-cDNA was then released from streptavidin beads, circularized and re-linearized, producing single-stranded DNA in the form of left.half.Linker-cDNA-DNA-right.half.Linker. The two halves of the linker (left.half.Linker and right.half.Linker) served as templates for PCR amplification. The linearized DNA was amplified with NEBNext PCR primers for Illumina, size-selected, and subjected to 100 cycles of pair-end sequencing with an Illumina Hi-seq 4000. Approximately 300 million read pairs were obtained per sample. The sequencing data were analyzed using a standard processing pipeline iMARGI-Docker (https://sysbio.ucsd.edu/imargi_pipeline).

Super enhancers (SEs) for HUVECs were downloaded from dbSUPER^30^. The 912 super enhancers were classified into three categories: not overlapping any gene (84), overlapping one or more genes but not fully embedded within any of them (449) and overlapping one or more genes and fully embedded within at least one of them (379). For these 379 SEs, when the SE is fully embedded into a gene, the entire gene was considered a SE. We thus updated the start and end coordinate per each super enhancer SE_i as following: we extracted all the genes SE_i is embedded within (SE_i_genes); then we replaced SE_i start coordinate with the minimum coordinate of SE_i_genes, and SE_i end coordinate with the maximum end coordinate of SE_i_genes. Given that some SEs could be embedded into the same genes or overlapping genes, it may happen that, after having updated the coordinates some of these SEs precisely overlap each other. After having removed these “duplicates”, we obtained 875 super enhancers that will be used in the following analysis. The sum of the lengths of these SEs is 94,493,925 bp (~108 kb on average per SE, 875 super enhancers in total). If we consider the entire genome length (3,088,286,401 bp), we have that 3.1% of the entire genome is occupied by super enhancers.

In order to analyze super enhancer interactions, only highly interacting super enhancers were considered, i.e. super enhancers with a high number of read pairs mapped over them. All the super enhancer pairs with a low number of reads mapped over them were discarded. This was achieved by setting a threshold *gamma* on the ratio between the read pairs among SEs and the total pairs in each sample (R_SE_total), to avoid any bias caused by the different sample sizes. The threshold was chosen as the 99th percentile of the distribution of R_SE_total for inter-chromosomal interacting SE pairs in control sample (Figure S4). The tendency of increase and decrease of interacting inter- and intra-chromosomal super enhancer pairs respectively, was independent of the choice of the threshold gamma (Table S4). SE hubs were then selected exploiting the degree of node (DON), i.e. the number of edges per each SE (node) of the networks. SEs with a DON > 60 were marked as hubs. This threshold was strictly selected as over the 95th percentile values of the DON distributions for all the three conditions (Figure S5).

### DNA-FISH experiment and imaging analysis

DNA FISH probes were obtained from Empire Genomics, LLC. and tested for hybridization specificity. Information of DNA FISH probe design is listed in Table SXX. Probes used for DNA FISH were designed against genomic regions overlapping LINC00607-SERPINE chimeric read aligned regions. Region of chromosome 2 that exhibited strong hybridization signal in FISH, but do not show significant interacting signal with SERPINE1 SE, was used as the control.

DNA-FISH was performed by following probe manufacturer protocol (Empire genomics) with minor modifications. Briefly, for imaging experiment, HUVEC cells were seeded onto Poly-L-Lysine pre-coated coverslips (Fisher Scientific, #NC0326897), and treatment was performed as described. All incubation steps were performed at RT unless otherwise indicated. On Day 3 and Day 7 of treatment, cells were washed twice in PBS and fixed in fresh 4% paraformaldehyde (pH 7.2) for 30 mins.

Cells were permeabilized in 0.1% saponin (Sigma-Aldrich, #84510), 0.1% Triton-X 100 in PBS for 10 mins at RT. Subsequently, cells were incubated in 20% glycerol in PBS for 20 mins, followed by three times of freeze-thaw in liquid nitrogen. The slides were then denatured in 0.1 M HCl for 30 mins, and blocked in 3% BSA and 100 ug/ml RNase A in PBS for 1 hr at 37 °C. This was followed by a second permeabilization step in 0.5% saponin/ 0.5 % triton-X 100 in PBS for 30 mins. The slides were then denatured sequentially in 70 % formamide (Invitrogen, #AM9342)/ 2X SSC (Sigma-Aldrich, #S6639) for 2.5 mins, then 50 % formamide/ 2X SSC for 1 min in 73 °C water bath, before immediately incubated with probe mixtures denatured at 75 °C for 5 mins. Probes were prepared according to manufacturer protocol. After 18 hrs of hybridization in a dark humid chamber, the slides were washed with agitation in the following solutions sequentially: 50 % formamide / 2X SSC, 2X SSC (37 °C for both) and 4X SSC / 0.1 % tween. DAPI-staining and slides-mounting (Thermo Fisher Scientific, #00-4958-02) were performed after PBS rinse.

Slides were imaged using Perkin Elmer UltraView Vox Spinning Disk Confocal ×63 oil-immersion objective lens. Distance between red and green fluorescent FISH signal spots in the nucleus was quantified using Fiji (http://fiji.sc/wiki/index.php/Fiji) and Matlab softwares. P-values were generated using nonparametric Wilcoxon test with Bonferroni correction for multiple comparisons. Statistical tests were performed in R.

### Subcellular fractionation and RNA isolation

Subcellular fractionation was performed as reported previously^8^ with minor modification. Briefly, HUVECs from three confluent 150 mm culture dishes were applied as independent triplicates. The cells were collected in 200 μl cold cytoplasmic lysis buffer (0.15% NP-40, 10 mM Tris pH 7.5, 150 mM NaCl) and incubated on ice for 5 mins. The lysate was layered onto 500 μl cold sucrose buffer (10 mM Tris pH 7.5, 150 mM NaCl, 24% sucrose weight by volume) and centrifuged. The supernatant containing cytoplasmic component was quickly added to TRizol LS for RNA extraction. The nuclear pellet was gently suspended into 200 μl cold glycerol buffer (20 mM Tris pH 7.9, 75 mM NaCl, 0.5 mM EDTA, 50% glycerol, 0.85 mM DTT). An addition of cold nuclei lysis buffer (20 mM HEPES pH 7.6, 7.5 mM MgCl_2_, 0.2 mM EDTA, 0.3 M NaCl, 1 M urea, 1% NP-40, 1 mM DTT) was added, followed by vortex and centrifuge. The supernatant containing nucleoplasmic fraction was mixed with TRizol LS for RNA extraction. Cold PBS (50 ml) was added to the remaining pellet and gently pipetted. After vigorous vortex to resuspend the chromatin, chromatin-associated RNA was extracted by adding 100 μl chloroform and TRizol reagent. RNA samples from three different fractions were dissolved with same amount of RNase-free water and same volume of RNA was used for reverse transcription and qPCR.

### Monocyte adhesion assay

Monocytes adhesion assay was performed as previously described^8^. CD14+-sorted human primary monocytes from 4 different healthy donors were labeled with CellTracker™ Green CMFDA Dye and incubated with monolayer ECs (4 × 10^3^cells cm^−2^) for 15-30 mins in a cell culture incubator. The non-attached monocytes were then washed off with complete EC growth medium. The attached monocyte numbers were evaluated on Cytation™ 1 Cell Imaging Multi-Mode Reader (BioTek) using green fluorescent channel. Average numbers per sample were calculated from five randomly selected fields.

### SA-β-gal staining

Cytochemical staining for SA-β-galactosidase was performed using the Senescence β-Galactosidase Staining Kit following the manufacturer’s manual. Briefly, the endothelial cells post transfection and H+T treatment were washed once with freshly-prepared 1X PBS, fixed in 1X Fixative solution for 10-15 mins at room temperature, and then rinsed twice with 1X PBS. The cells were stained for 48 hours in a dry incubator before viewing under a Leica DMi1 microscope (Leica microsystems) at 50x magnifications. The percentage of SA-β-galactosidase positive cells was determined by counting the number of blue cells under bright field illumination.

### Statistical analysis

For all experiments in Figures 1 and 7, at least three independent experiments were performed unless otherwise specified. Statistical analysis was performed using Student’s *t*-test (two-sided) between two groups or ANOVA followed by Bonferroni post-test for multiple-group comparisons. *p*<0.05 was considered as statistically significant. For other experiments, the quantification and statistical analysis have been specified and detailed in figure legends and methods.

## Data availability

All data supporting the current study are available in the article and its Supplemental Information Files or are available from the corresponding authors on request.

## Acknowledgments

This work was supported by NIH grants R00HL122368 and R01HL121365 (to Z.B.C.) and DP1HD087990 and NIH 4D Nucleome U01CA200147 (to S.Z.), and Ella Fitzgerald Foundation (to Z.B.C.). Research reported in this publication included work performed in the Integrative Genomics Core at City of Hope supported by the National Cancer Institute of the National Institutes of Health under award number P30CA033572.

## Author Contributions

Conceptualization, Z.B.C. and S.Z.; Methodology, Z.B.C., S.Z., R.C., L.X., Y.L., X.F., C.C., and T.N; Investigation and validation, L.X., Y.L., R.C., K.S., X.F., C.C., and T.N., writing-original draft, Z.B.C, R.C., L.X., S.Z.; Writing-review and editing, Z.B.C, S.Z., R.C., R.N., and Y.L.; Resources, Z.B.C, S.Z., and R.N., Supervision, Z.B.C and S.Z.; Funding acquisition, Z.B.C and S.Z..

## Competing Financial Interest Statement

N/A

