## Supplemental Information Files for "Dynamic changes in RNA-chromatin interactome promote endothelial dysfunction"

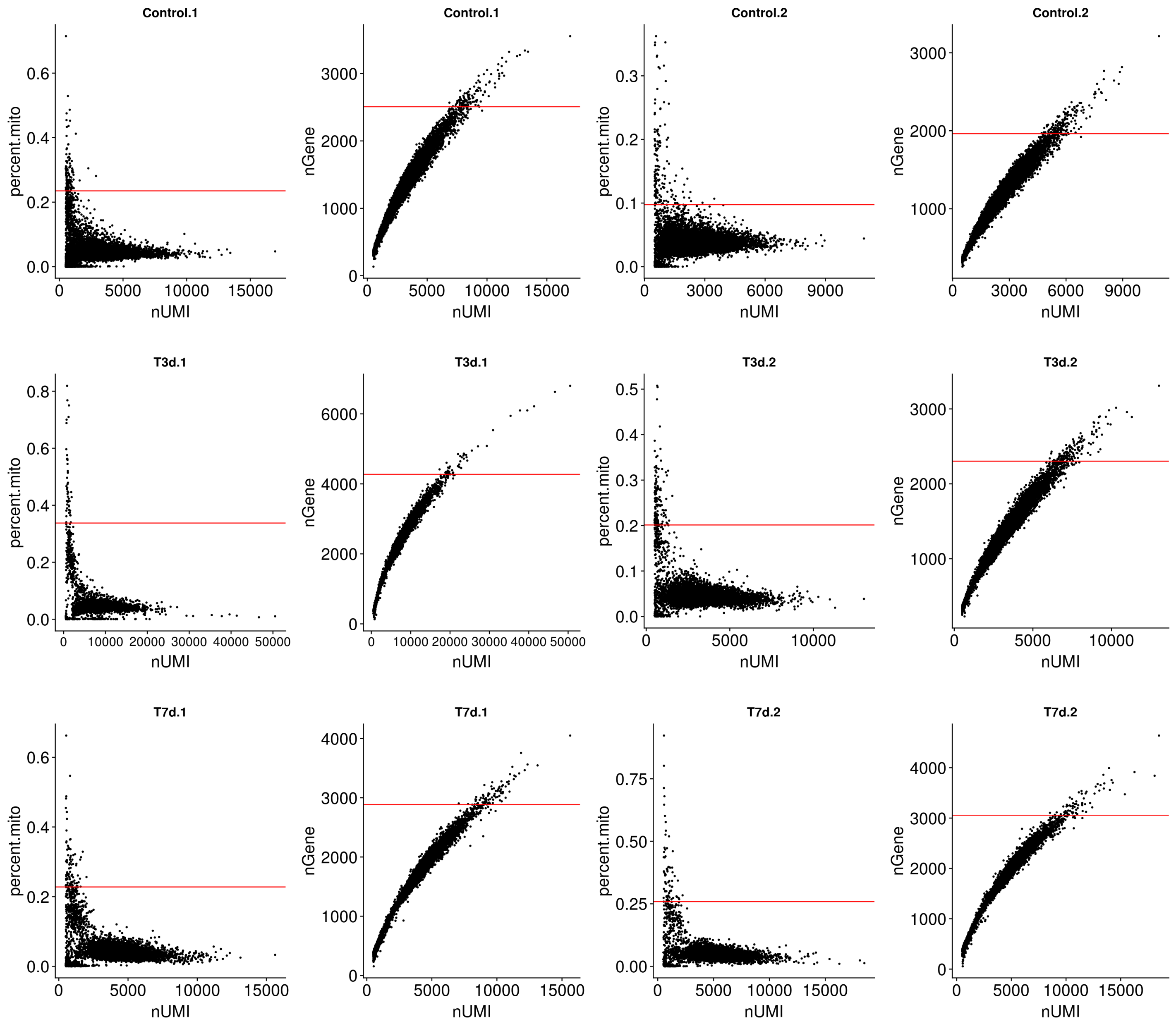


**Figure S1. Scatter Plots of UMI Counts (Numi) with Mitochondrial Percentage (Percent.Mito) and Gene Counts (Ngene) per cell in Each Sample, Related to Figure 2**

Red lines represent the thresholds used to filter out cells with outlier levels of mitochondrial percentage and gene counts (99$^{th}$ percentile).

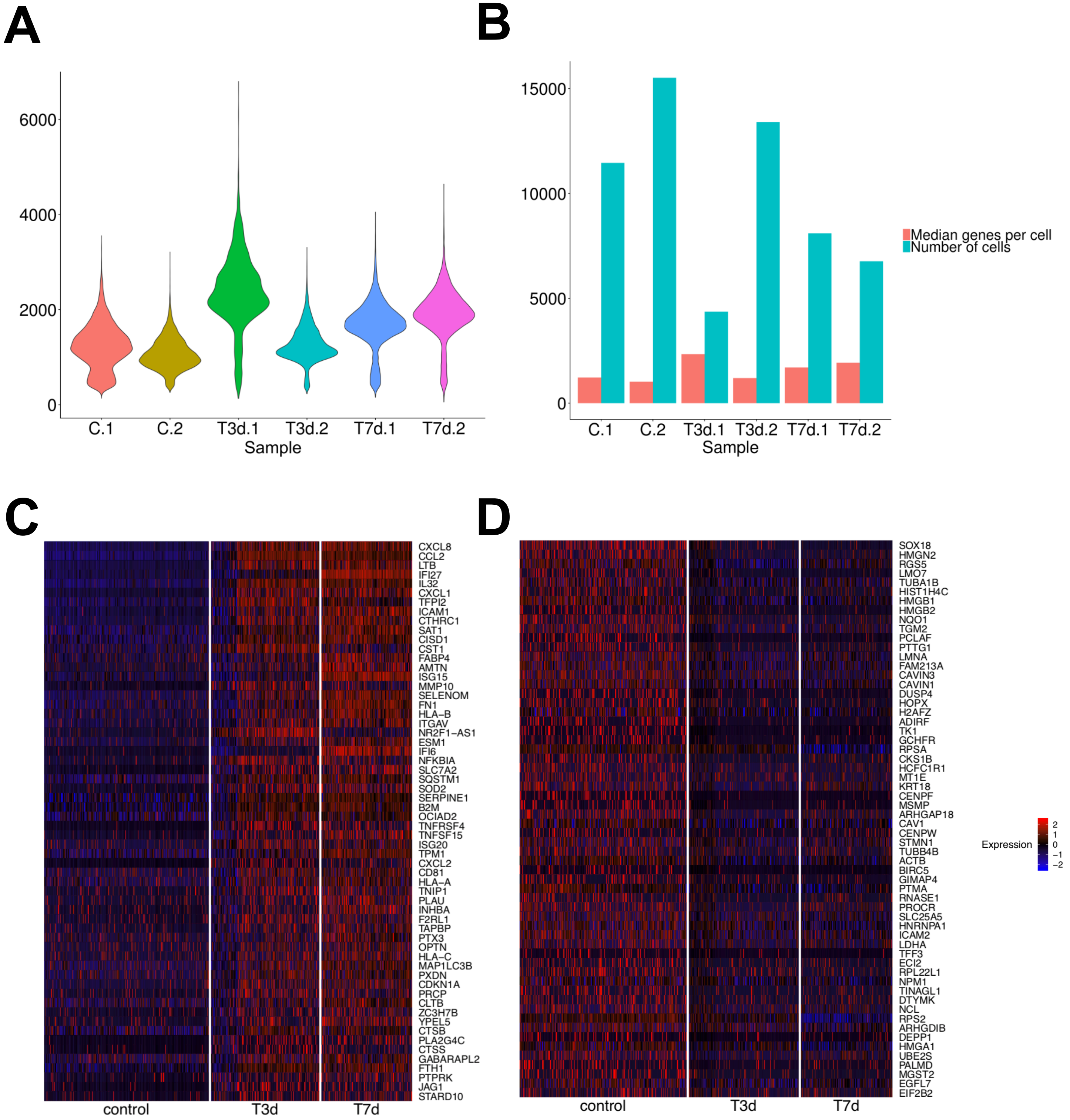


**Figure S2. Single-cell RNA-seq Data Quality Control and Heat Maps of Differential Expressed Genes, Related to Figure 2.**

(A) Violin plots of numbers of genes per single cell in each sample.

(B) Median genes per single cell and number of cells sequenced in each sample.

(C) Heatmap of gene expression of top 60 consistently up-regulated genes sorted by the mean value between the two avg_logFC (the one between Day 3 and control and the one between Day 7 and control)

(D) Heatmap of gene expression of top 60 consistently down-regulated genes sorted by the mean value between the two avg_logFC (the one between Day 3 and control and the one between Day 7 and control)

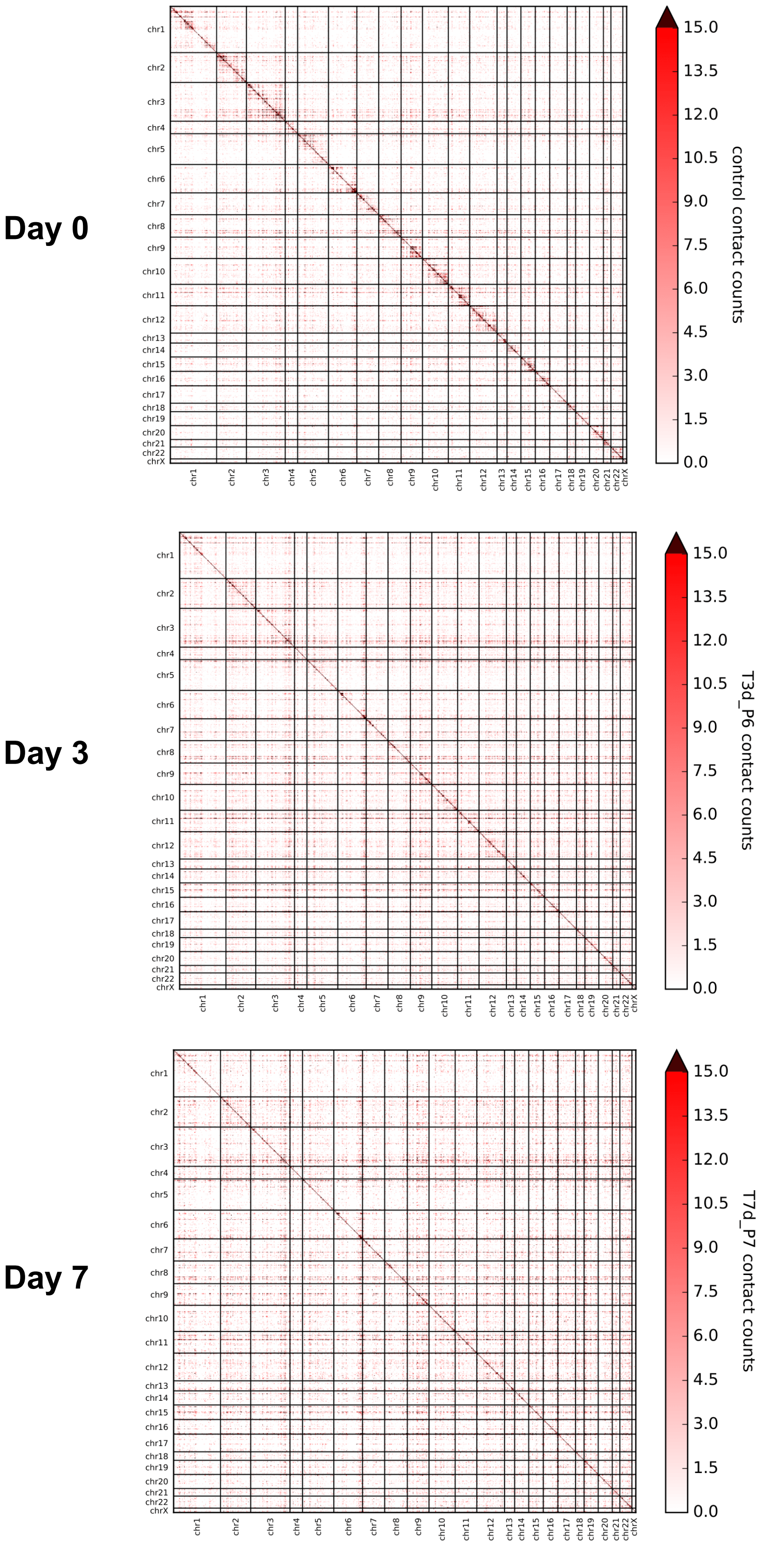


**Figure S3. iMARGI profiled-global heat maps for SE, Related to Figure 3.**
Each bin represents a SE; each value is the number of read pairs between SE normalized by the total number of read pairs per sample.

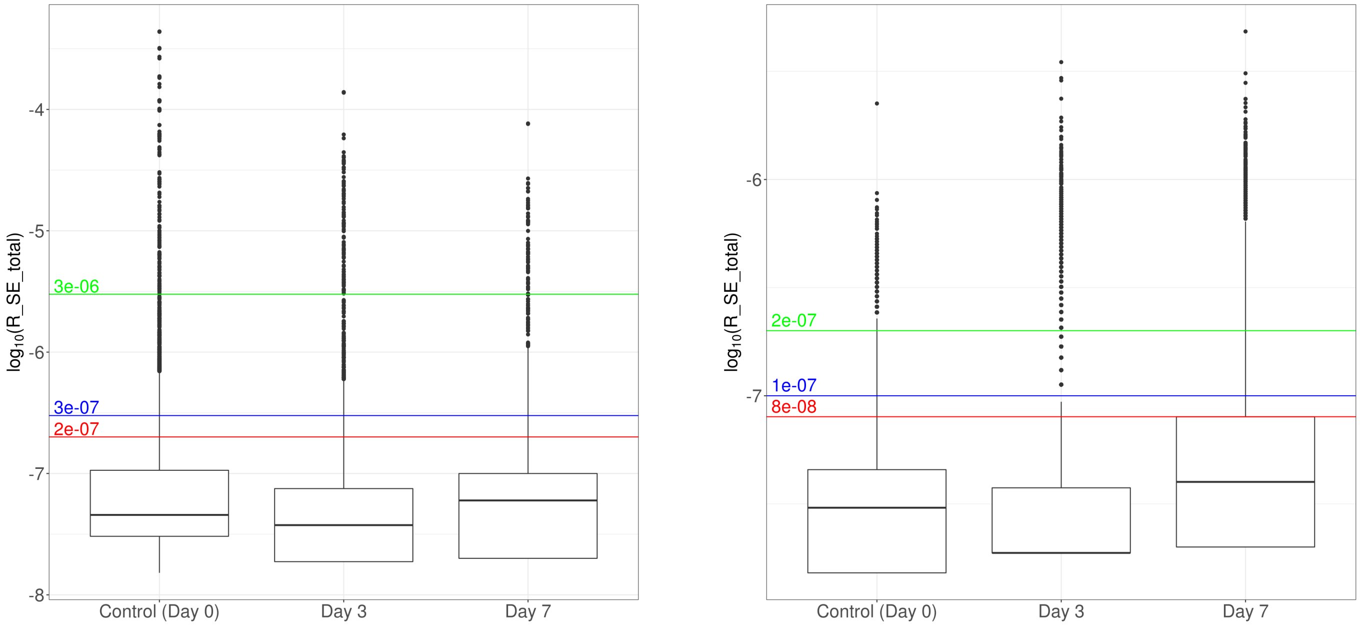


**Figure S4. Boxplot of the log10(R_SE_total) distribution for intra-chromosomal (left) and inter-chromosomal (right) SE interacting pairs, Related to Figure 5.**

Three thresholds are plotted with different colors: 90th percentile (red), 95th percentile (blue) and 99th percentile (green). R_SE_total: ratio between the number of iMARGI read pairs mapped over each super enhancer pair and the total read pairs in each sample.

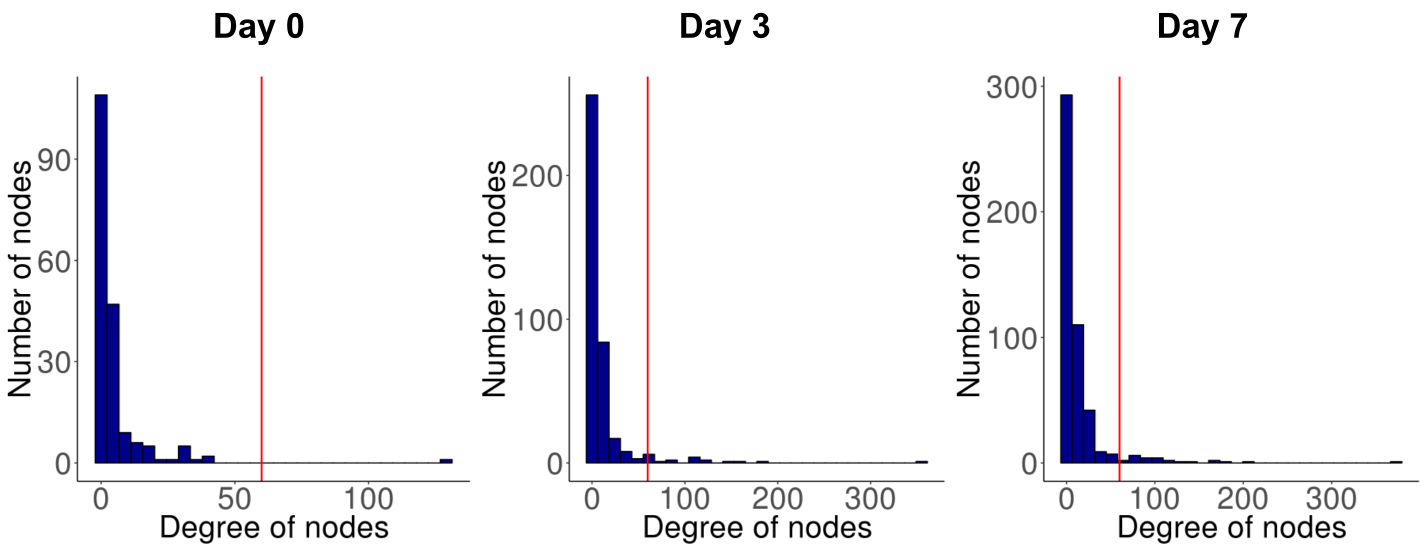


**Figure S5. Distribution of degree of nodes (i.e. the number of edges incident to that node) for three conditions, Related to Figure 5.**
The red lines represent DON values at 60, which are over the 95th percentile of the DON distribution for all three conditions.

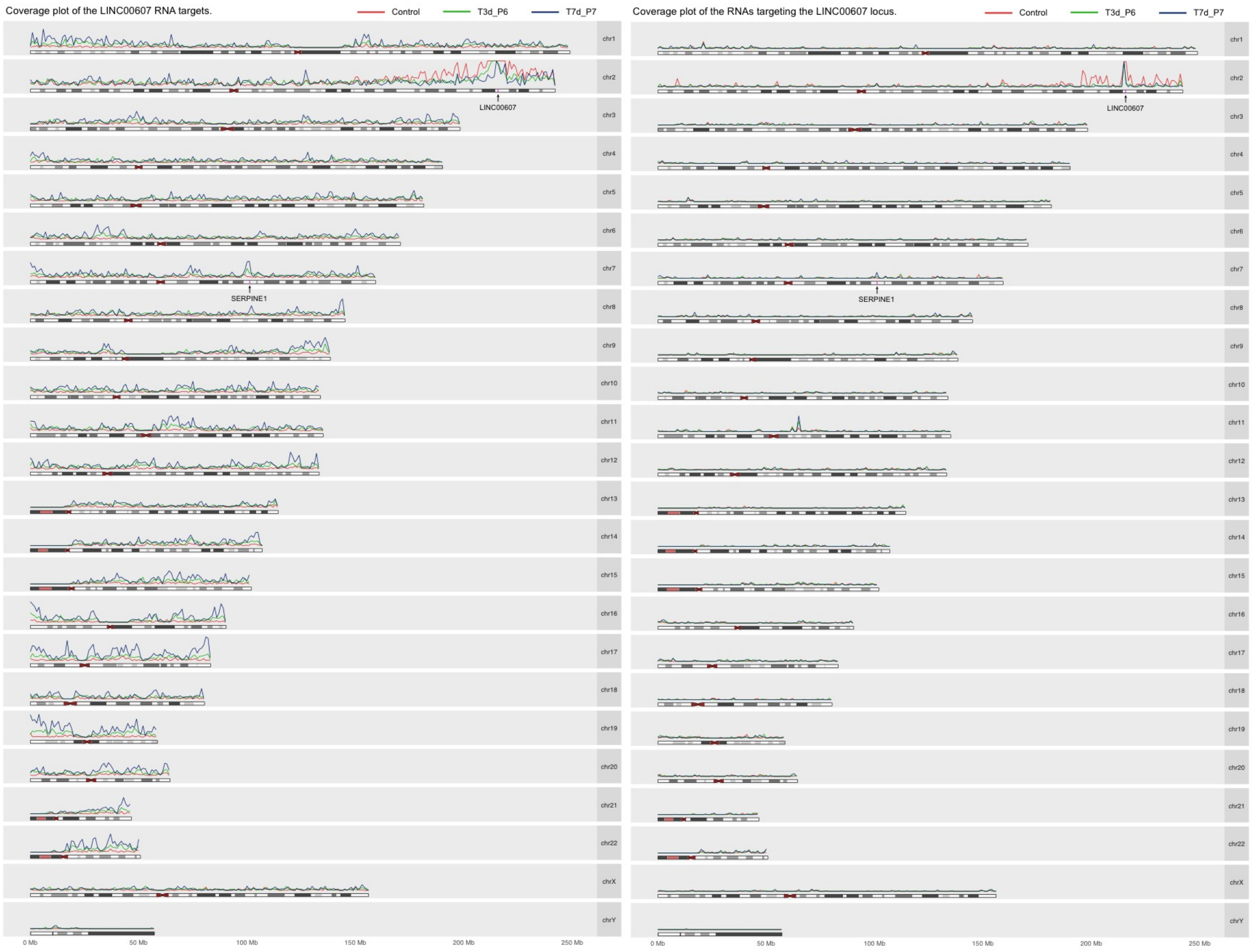


**Figure S6. iMARGI mapped LINC00607-SERPINE1 interaction, Related to Figure 6.**
Left panel is the coverage plot of the LINC00607 RNA targets. Right panel is the coverage plot of the RNAs targeting the LINC00607 locus.

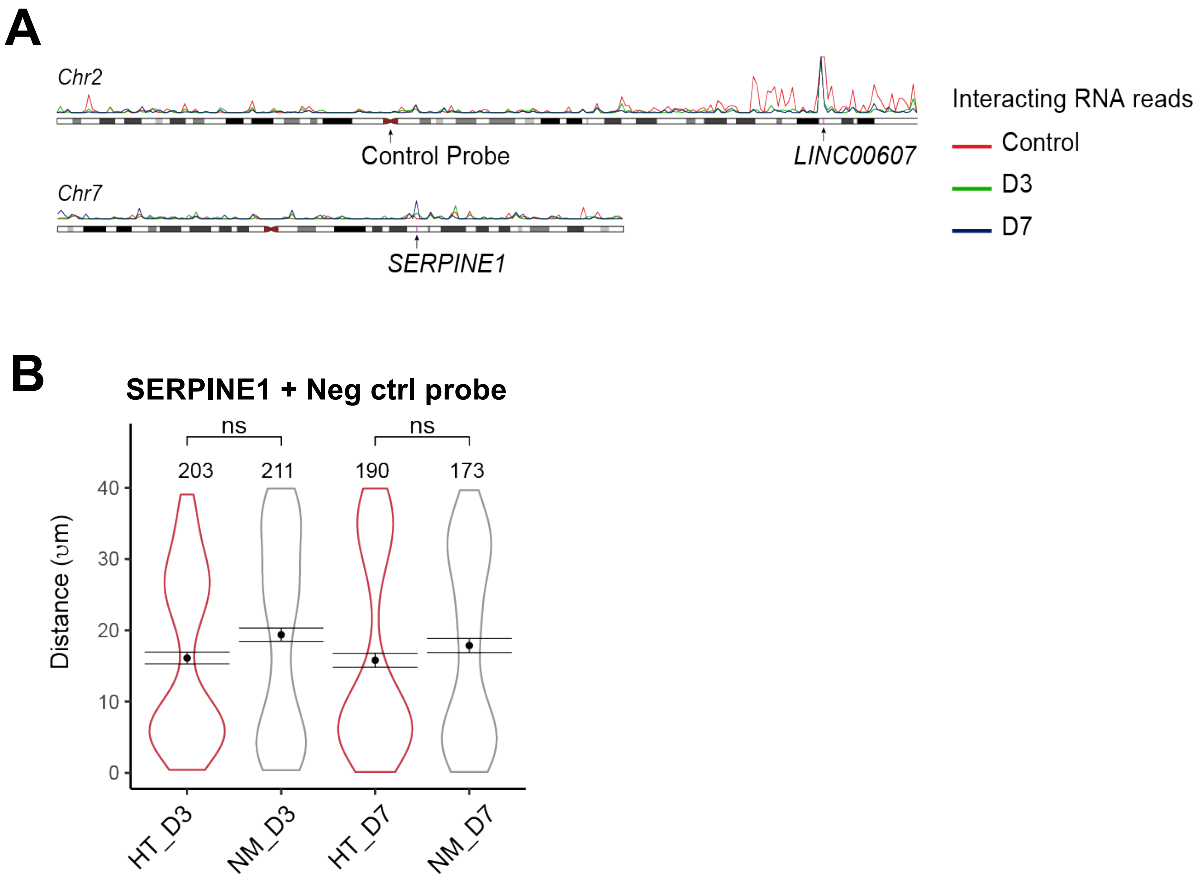


**Figure S7. Design of DNA FISH probes and Distance between SERPINE1 and control regions, Related to Figure 6.**

(A) Design of probes targeting chromosomal loci spanning *LINC00607* and *SERPINE1* enhancers, as well as the negative control region.

(B) Distances between SERPINE1 genomic locus and the control region were measured on Day 3 or Day 7 of H+T treatment (HT) or normal glucose and osmolarity control (NM). Quantification was based on FISH signal from probes for *SERPINE1* and control region. Total numbers of pairs are listed on top of the plot. p-values were generated by nonparametric Wilcox-test with Bonferroni correction for multiple comparisons.

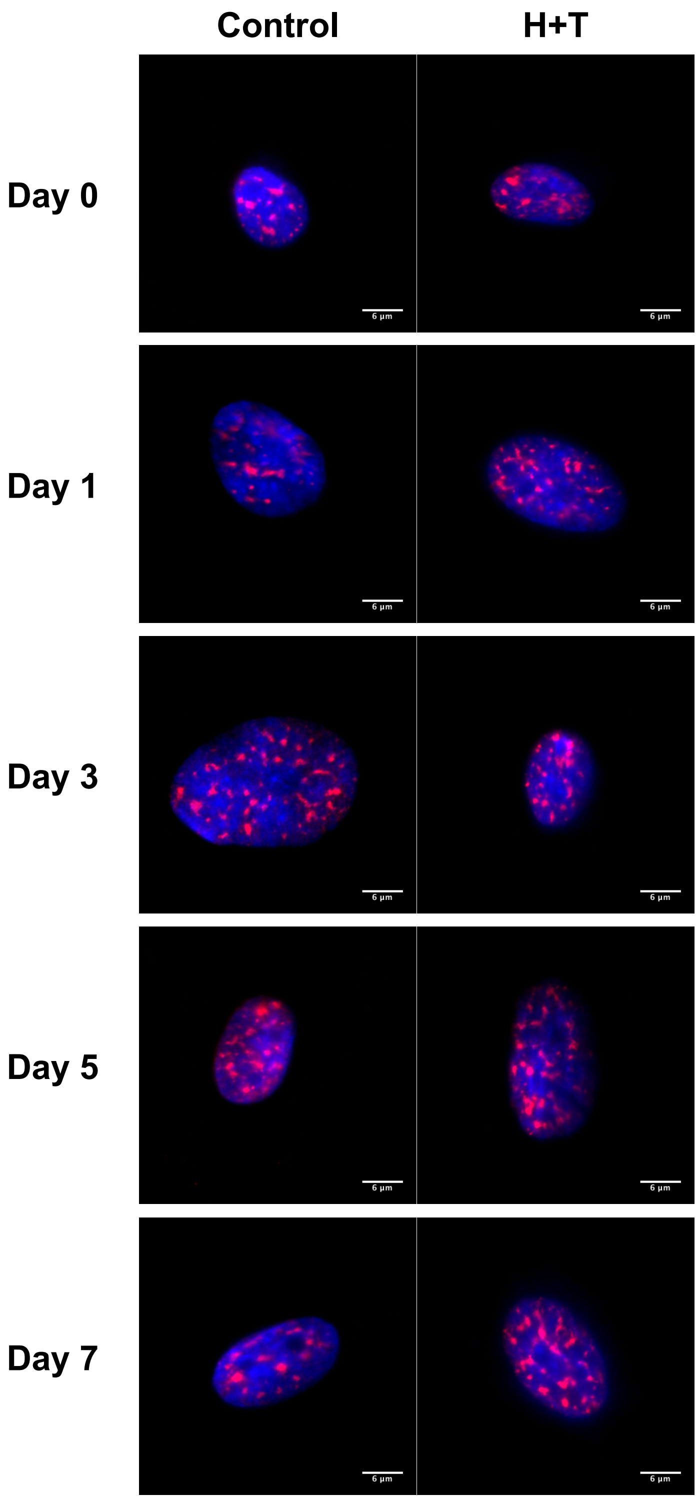


**Figure S8. IF staining of SC35 in HUVECs, Related to Discussion.**
HUVECs were stained with anti-SC35 (red). The nuclei were imaged with DAPI (blue). Each image represents the 30th z-stack of each sample.

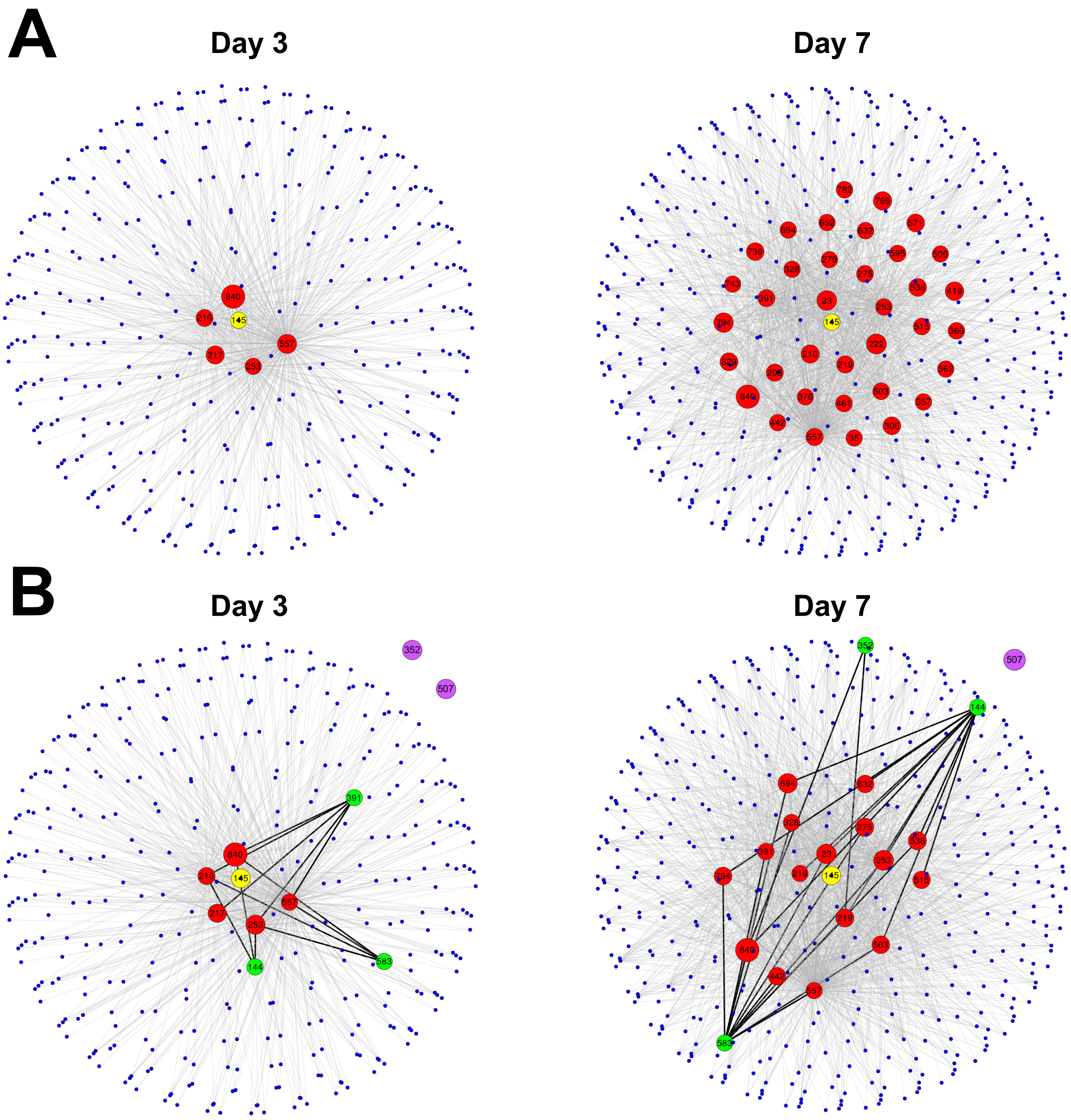


**Figure S9. Two-step network of LINC00607 interacting SEs, Related to Discussion.**
(A) Two-step network with LINC00607 (node in yellow) as the origin. LINC00607 does not appear at all in the control sample (not plotted), while the network dimension increased from Day 3 to Day 7. Red nodes represent SEs directly interacting with LINC00607.
(B) Two-step networks highlighting the SEs directly interacting with LINC00607 and also interacting with markers whose expression were suppressed by LINC00607 knockdown (e.g., CTGF, SERPINE1, and FN1). Markers within the two-step network are plotted in green, whereas those plotted in purple are standalone nodes. Edges between nodes directly interacting with LINC00607 and the select markers are plotted thicker.

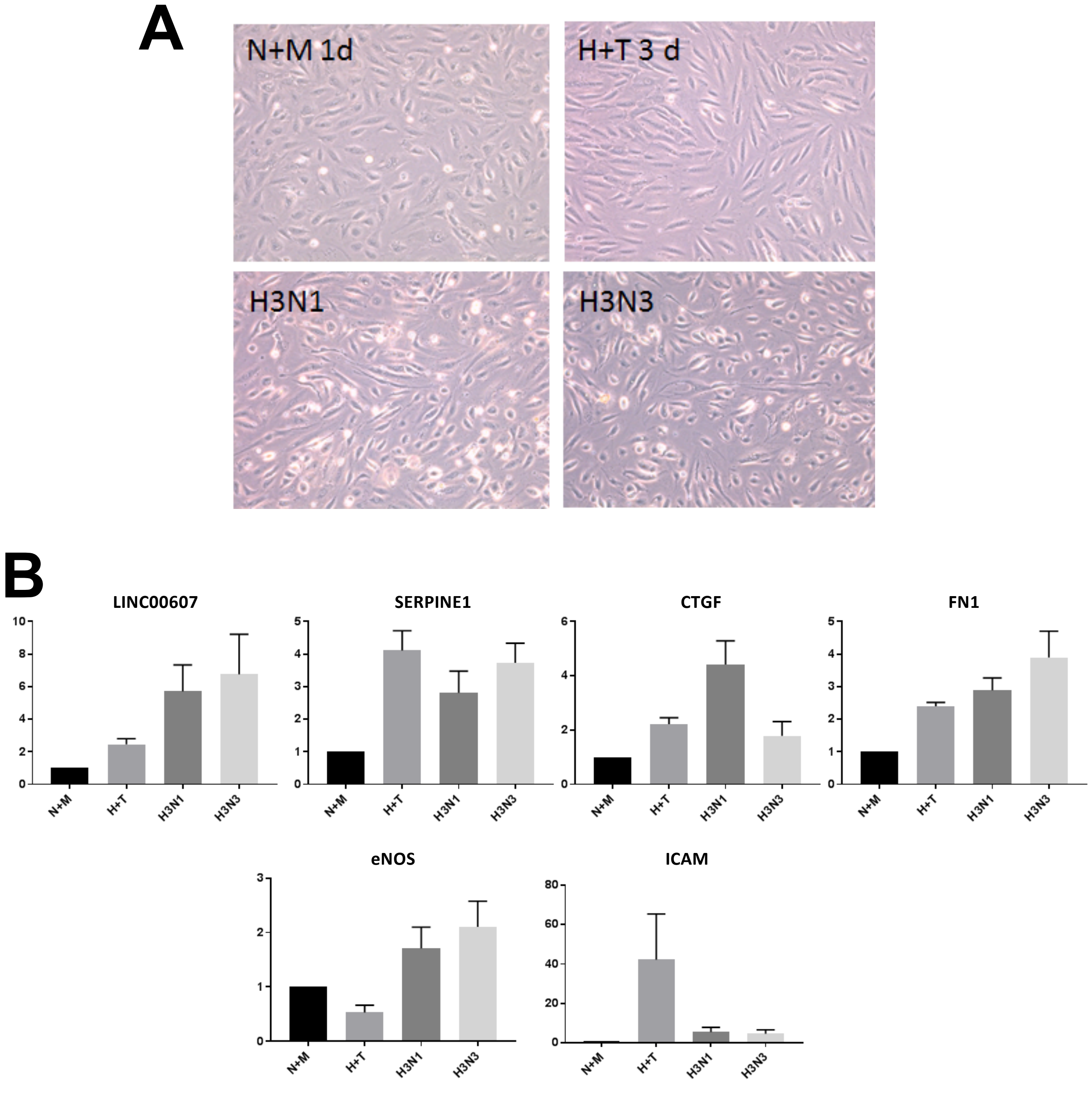


**Figure S10. Partial reversal effect of removal of H+T stimuli, Related to Discussion.**
(A) Morphological changes of HUVECs upon mannitol (N+M 1d) or H+T treatment for 3 days (H+T 3d) followed by incubation in mannitol for 1 day (H3N1) or 3 days (H3N3).
(B) Relative RNA levels of indicated mRNAs or LINC00607 under various treatments as shown in (A).

**Table S1. Enriched pathways in DE genes related to EC dysfunction, Related to Figure 2.**

| Term | Count | % | PValue |
|---|---|---|---|
| GO:0007155~cell adhesion | 43 | 5.34825871 | 3.32E-06 |
| GO:0001525~angiogenesis | 37 | 4.60199005 | 6.02E-12 |
| GO:0006954~inflammatory response | 35 | 4.35323383 | 4.24E-05 |
| GO:0006955~immune response | 32 | 3.9800995 | 0.00252838 |
| GO:0098609~cell-cell adhesion | 31 | 3.85572139 | 2.04E-06 |
| GO:0001666~response to hypoxia | 26 | 3.23383085 | 8.91E-08 |
| GO:0030198~extracellular matrix organization | 23 | 2.86069652 | 3.73E-05 |
| GO:0016477~cell migration | 20 | 2.48756219 | 1.53E-04 |
| GO:0050900~leukocyte migration | 17 | 2.11442786 | 6.66E-05 |
| GO:0007568~aging | 16 | 1.99004975 | 0.00501572 |
| GO:0007179~transforming growth factor beta receptor signaling pathway | 15 | 1.86567164 | 3.60E-05 |
| GO:0033209~tumor necrosis factor-mediated signaling pathway | 14 | 1.74129353 | 0.00166058 |
| GO:0006979~response to oxidative stress | 13 | 1.61691542 | 0.0026811 |
| GO:0050776~regulation of immune response | 13 | 1.61691542 | 0.08217528 |
| GO:0008286~insulin receptor signaling pathway | 12 | 1.49253731 | 4.78E-04 |
| GO:0071260~cellular response to mechanical stimulus | 11 | 1.3681592 | 8.64E-04 |
| GO:0034097~response to cytokine | 10 | 1.24378109 | 3.31E-04 |
| GO:0071356~cellular response to tumor necrosis factor | 10 | 1.24378109 | 0.04704454 |
| GO:0007219~Notch signaling pathway | 10 | 1.24378109 | 0.05925478 |
| GO:0009749~response to glucose | 9 | 1.11940299 | 0.00846146 |
| GO:0050727~regulation of inflammatory response | 8 | 0.99502488 | 0.01797915 |
| GO:0050729~positive regulation of inflammatory response | 8 | 0.99502488 | 0.03683555 |
| GO:0032869~cellular response to insulin stimulus | 8 | 0.99502488 | 0.04706729 |
| GO:0045429~positive regulation of nitric oxide biosynthetic process | 7 | 0.87064677 | 0.00965174 |
| GO:0071560~cellular response to transforming growth factor beta stimulus | 7 | 0.87064677 | 0.01784758 |
| GO:0007249~I-kappaB kinase/NF-kappaB signaling | 7 | 0.87064677 | 0.04320139 |
| GO:0038061~NIK/NF-kappaB signaling | 7 | 0.87064677 | 0.06346437 |
| GO:0006006~glucose metabolic process | 7 | 0.87064677 | 0.06730356 |
| GO:0070098~chemokine-mediated signaling pathway | 7 | 0.87064677 | 0.08397872 |
| GO:0090023~positive regulation of neutrophil chemotaxis | 6 | 0.74626866 | 0.00206053 |
| GO:0001974~blood vessel remodeling | 5 | 0.62189055 | 0.04675183 |

**Table S2. Overview and Distribution of HiC Data, Related to Figure 3.**

| | Control (Day 0) | Day 3 | Day 7 |
|---|---|---|---|
| **Total read pairs** | 186,661,191 | 200,746,376 | 106,348,319 |
| **Intra-chromosomal read pairs** | 172,897,716 (92.6%) | 183,479,138 (91.4%) | 99,736,000 (93.8%) |
| **Inter-chromosomal read pairs** | 13,763,475 (7.4%) | 17,267,238 (8.6%) | 6,612,319 (6.2%) |

**Table S3. Overall read pairs mapped by iMARGI across three conditions, Related to Figure 4.**

| | Control (Day 0) | Day 3 | Day 7 |
|---|---|---|---|
| **Total read pairs** | 65,879,147 | 53,291,562 | 50,031,654 |
| **Intra-chromosomal read pairs** | 45,041,984 (68.4%) | 22,502,605 (42.2%) | 14,858,759 (29.7%) |
| **Inter-chromosomal read pairs** | 20,837,163 (31.6%) | 30,788,957 (57.8%) | 35,172,895 (70.3%) |
| **Read pairs with RNA end over SEs** | 10,574,927 (16%) | 7,691,827 (14.4%) | 7,206,673 (14.4%) |
| **Read pairs with DNA end over SEs** | 6,316,504 (9.6%) | 2,967,038 (5.6%) | 2,532,498 (5.1%) |
| **Read pairs both over SEs** | 4,729,619 (7.2%) | 1,506,712 (2.8%) | 931,370 (1.9%) |
| **Intrachromosomal read pairs within SEs** | 100,331 | 38,896 | 27,691 |
| **Intrachromosomal read pairs within SEs per total read pairs** | 1.5e-3 | 0.73e-3 | 0.55e-3 |
| **Interchromosomal read pairs within SEs** | 179,283 | 246,086 | 260,659 |
| **Interchromosomal read pairs within SEs per total read pairs** | 2.7e-3 | 4.6e-3 | 5.2e-3 |

**Table S4. Numbers of SE interacting pairs above different thresholds across three conditions. Number of interacting super enhancer pairs (edges) and super enhancers (nodes, between parentheses) in intra- and inter-chromosomal networks at different thresholds gamma (between parentheses the correspondent percentile per each threshold). The fold change (FC) between the number of interacting super enhancer pairs in Day 7 and control sample is also reported, Related in Figure 4.**

| | ***gamma*** | **Control (Day 0)** | **Day 3** | **Day 7** | **FC (Day 7 vs control)** |
|---|---|---|---|---|---|
| **Intra-chromosomal networks** | 2e-7 (90th) | 1,609 (596) | 789 (462) | 621 (398) | -2.6 |
| | 3e-7 (95th) | 1,119 (515) | 474 (359) | 375 (303) | -3 |
| | 3e-6 (99th) | 141 (147) | 96 (98) | 69 (70) | -2 |
| **Inter-chromosomal networks** | 8e-8 (90th) | 5,784 (621) | 10,910 (655) | 17,275 (825) | 3 |
| | 1e-7 (95th) | 3,903 (529) | 7,570 (588) | 12,171 (782) | 3.1 |
| | 2e-7 (95th) | 510 (187) | 2,234 (387) | 3,367 (486) | 6.6 |

**Table S5. List of hubs (SE indexes sorted numerically), Related to Figure 6.**

| SE_index | SE_genes |
|---|---|
| 23 | MACF1;RNU6-608P;RNA5SP44;RP11-416A14.1;RP11-420K8.1;HSPE1P8 |
| 216 | FNDC3B;RP11-423E7.1;RN7SL141P;RP11-423E7.2 |
| 217 | NLGN1;RP11-521A24.1;NLGN1-AS1;RN7SKP234 |
| 219 | RP11-778D9.12;EIF2B5;RP11-778D9.4;DVL3;AP2M1;ABCF3;VWA5B2;MIR1224;ALG3;ECE2;CAMK2N2;PSMD2;EIF4G1;SNORD66;FAM131A;CLCN2;POLR2H;THPO;CHRD;RP11-433C9.2;EIF2B5-IT1;EIF2B5-AS1;EPHB3 |
| 253 | TRIO;AC016549.1;AC016656.1 |
| 264 | PDE4D;RP11-266N13.2;Clostridiales-1;CTD-2146O16.1;NDUFB4P2;MIR582;CTD-2254N19.1;RNU6-806P;RP11-546M4.1;AC109486.1;SETP21;PART1_1;PART1;PART1_2 |
| 328 | CASC15;RP4-551N11.1;RN7SKP240;NBAT1;RP11-524C21.1 |
| 391 | TRIM56;SERPINE1 |
| 438 | EXT1 |
| 442 | PVT1;PVT1_1;MIR1204;PVT1_3;TMEM75;MIR1205;RNU1-106P;RP11-55J15.2;RNU4-25P;MIR1207 |
| 469 | PALM2;RP11-406O23.2;PALM2-AKAP2;RP11-151F5.2;AKAP2 |
| 470 | PALM2;PALM2-AKAP2;RP11-151F5.2;AKAP2 |
| 503 | RP11-295P9.13;RP11-295P9.3;FRMD4A;RP11-295P9.12;AL157392.1;RP11-295P9.6;RNA5SP301;RP11-353M9.1;NUTF2P5;RP11-142M10.2;RP11-397C18.2;MIR4293;Metazoa_SRP;MIR1265 |
| 548 | NAV2;NAV2-IT1;RNA5SP335;NAV2-AS5;NAV2-AS4;MIR4486;SNORA1;RP11-359E10.1;MIR4694;NAV2-AS3;NAV2-AS2;NAV2-AS1 |
| 556 | NEAT1;NEAT1_1;NEAT1_2;NEAT1_3 |
| 557 | CMB9-22P13.2;MALAT1;AP000769.7;AP000769.1;MALAT1;mascRNA-menRNA |
| 583 | VWF;RN7SL69P;RP3-454B23.1;CD9;Y_RNA |
| 632 | NCOR2;MIR6880;RP11-408I18.9;RP11-83B20.1;RP11-83B20.2;RP11-83B20.3;RP11-83B20.4;RP11-83B20.5;RP11-83B20.6 |
| 658 | SAMD4A;AL138994.1 |
| 682 | THBS1;CTD-2033D15.2;CTD-2033D15.3;CTD-2033D15.1 |
| 692 | SMAD3;RP11-342M21.2 |
| 694 | THSD4;CT62;THSD4-AS1;RP11-673C5.1;RP11-592N21.1;RP11-592N21.2;RP11-1123I8.1;AC104938.1 |
| 733 | ANKRD11;AC137932.4;AC137932.5;AC137932.6;RP1-168P16.2;RP1-168P16.1;RNU6-430P;RP1-168P16.3 |
| 794 | SIPA1L3;CTC-244M17.1;AC011465.1;CTC-450M9.1;CTB-102L5.7;CTB-102L5.8;RN7SL663P |
| 840 | RUNX1;AF015262.2;RPL34P3;EZH2P1;AF015720.3;MIR802;RPS20P1;PPP1R2P2;AP000687.1 |

**Table S6. List of SEs directly interacting with LINC00607 or select markers promoting EndoMT, Related to Discussion and Figure S9.**

| SE_index | SE_genes |
|---|---|
| 23 | MACF1;RNU6-608P;RNA5SP44;RP11-416A14.1;RP11-420K8.1;HSPE1P8 |
| 35 | PDE4B;RP11-397C12.1;RNU4-88P |
| 144 | FN1;AC012462.1 |
| 205 | WWTR1;RNU6-1098P;WWTR1-IT1;WWTR1-AS1 |
| 210 | MECOM;RP11-641D5.2;RP11-641D5.1;AC074033.1;RP11-3K16.2 |
| 216 | FNDC3B;RP11-423E7.1;RN7SL141P;RP11-423E7.2 |
| 217 | NLGN1;RP11-521A24.1;NLGN1-AS1;RN7SKP234 |
| 219 | RP11-778D9.12;EIF2B5;RP11-778D9.4;DVL3;AP2M1;ABCF3;VWA5B2;MIR1224;ALG3;ECE2;CAMK2N2;PSMD2;EIF4G1;SNORD66;FAM131A;CLCN2;POLR2H;THPO;CHRD;RP11-433C9.2;EIF2B5-IT1;EIF2B5-AS1;EPHB3 |
| 222 | LPP-AS2;LPP;FLJ42393;AC117507.1;LPP-AS1;AC063932.1;MIR28 |
| 253 | TRIO;AC016549.1;AC016656.1 |
| 275 | LHFPL2;HMGB1P21;CTD-2073O6.1;CTD-2045M21.1 |
| 279 | ELL2;CTD-2337A12.1;AC008592.7;FABP5P5;RP11-254I22.1;MIR583;RNU6-524P;RP11-254I22.3;RP11-254I22.2;PCSK1;CAST;AC020900.1;CTC-506B8.1;ERAP1 |
| 306 | DOCK2;CTB-37A13.1;FAM196B;AC008449.1;MIR378E |
| 328 | CASC15;RP4-551N11.1;RN7SKP240;NBAT1;RP11-524C21.1 |
| 352 | CTGF;RP11-69I8.3 |
| 357 | PDE7B;AL360178.1;AL512290.1;RP13-143G15.4;COX5BP2;RP11-472E5.3 |
| 362 | SCAF8;TIAM2;RP3-472M2.2;MIR1273C;AL121952.1;U8;RP1-66H9.1;RP11-477D19.2;TFB1M |
| 366 | PRKAR1B;AC147651.2;AC147651.5;AC147651.4;DNAAF5 |
| 376 | MALSU1;IGF2BP3;AC021876.4;SNORD65;RNU7-143P |
| 391 | TRIM56;SERPINE1 |
| 419 | TNFRSF10D |
| 442 | PVT1;PVT1_1;MIR1204;PVT1_3;TMEM75;MIR1205;RNU1-106P;RP11-55J15.2;RNU4-25P;MIR1207 |
| 461 | RP11-613M10.8;TOMM5;RP11-613M10.9;RAB1C;FRMPD1;RN7SKP171;TRMT10B;EXOSC3;DCAF10;RP11-3J10.7;PAICSP1;SLC25A51;TMX2P1;SHB;RNU7-124P |
| 503 | RP11-295P9.13;RP11-295P9.3;FRMD4A;RP11-295P9.12;AL157392.1;RP11-295P9.6;RNA5SP301;RP11-353M9.1;NUTF2P5;RP11-142M10.2;RP11-397C18.2;MIR4293;Metazoa_SRP;MIR1265 |
| 506 | CUBN |
| 507 | TRDMT1;VIM-AS1;VIM;RP11-124N14.3 |
| 515 | CCNY;RNU6-1167P;RP11-324I22.4 |
| 538 | ZRANB1;CTBP2;RP11-59C5.3;MIR4296 |
| 557 | CMB9-22P13.2;MALAT1;AP000769.7;AP000769.1;MALAT1;mascRNA-menRNA |
| 571 | DDX10;RPS2P39;RP11-801G16.2;CYCSP29 |
| 583 | VWF;RN7SL69P;RP3-454B23.1;CD9;Y_RNA |
| 595 | RP11-513G19.1;RP11-612B6.2;ITPR2;RNA5SP354;RP11-666F17.1;RP11-666F17.2 |
| 632 | NCOR2;MIR6880;RP11-408I18.9;RP11-83B20.1;RP11-83B20.2;RP11-83B20.3;RP11-83B20.4;RP11-83B20.5;RP11-83B20.6 |
| 652 | COL4A1;COL4A2;MIR8073;RP11-291I6.2;COL4A2-AS2;COL4A2-AS1 |
| 692 | SMAD3;RP11-342M21.2 |
| 694 | THSD4;CT62;THSD4-AS1;RP11-673C5.1;RP11-592N21.1;RP11-592N21.2;RP11-1123I8.1;AC104938.1 |
| 739 | RAI1;RAI1-AS1;SMCR5;RP1-253P7.1;SREBF1 |
| 743 | ARHGAP23;AC124789.1 |
| 769 | DLGAP1;RN7SL39P;RP11-710M11.1;DLGAP1-AS1;DLGAP1-AS2;RP11-874J12.4;RP11-874J12.3;DLGAP1-AS3;MIR6718;RNU6-831P;DLGAP1-AS4;GAPDHP66;DLGAP1-AS5;RP11-138C24.2 |
| 782 | GNG7;TCEB1P28;RN7SL121P;CTC-265F19.2;MIR7850;CTC-265F19.3;CTC-265F19.1;AC006538.8 |
| 794 | SIPA1L3;CTC-244M17.1;AC011465.1;CTC-450M9.1;CTB-102L5.7;CTB-102L5.8;RN7SL663P |
| 829 | COX6CP2;PTPN1;RN7SL672P;Y_RNA;RP4-530I15.9 |
| 840 | RUNX1;AF015262.2;RPL34P3;EZH2P1;AF015720.3;MIR802;RPS20P1;PPP1R2P2;AP000687.1 |

**Table S7. Sequences of qPCR primers and LNA Gapmers**

| qPCR primers | Forward | Reverse |
|---|---|---|
| LINC00607 | ACCGGGCGTTGAGAATACAA | ACACTTGGCGAAACTTCCCT |
| eNOS | TGATGGCGAAGCGAGTGAAG | ACTCATCCATACACAGGACCC |
| ICAM1 | GTGTCCTGTATGGCCCCCGACT | ACCTTGCGGGTGACCTCCCC |
| ACTB | CATGTACGTTGCTATCCAGGC | CTCCTTAATGTCACGCACGAT |
| COL4A1 | GGACTACCTGGAACAAAAGGG | GCCAAGTATCTCACCTGGATCA |
| FN1 | ACTGTACATGCTTCGGTCAG | AGTCTCTGAATCCTGGCATTG |
| CTGF | TAGGCTTGGAGATTTTGGGA | GGTTACCAATGACAACGCCT |
| SERPINE1 | AGTGGACTTTTCAGAGGTGGA | GCCGTTGAAGTAGAGGGCATT |
| α-SMA | CAGGGCTGTTTTCCCATCCAT | GCCATGTTCTATCGGGTACTTC |
| SMAD3 | CCATCTCCTACTACGAGCTGAA | CACTGCTGCATTCCTGTTGAC |
| **LNAs** | **Sequence** | **Position (NR_037195.1)** |
| LNA-1 | ATAGGTCACGCATTCT | 210-225 |
| LNA-2 | CAACTGTGGAATGATA | 2033-2048 |
| Scramble | AACACGTCTATACGC | |

**Information of DNA FISH probe**

| SKU | FISH Probes | Chromosome coordinate (Hg38) |
|---|---|---|
| CLN-1008 | Clone Library: RPCI-11<br>Clone Name: 946O2<br>Dye Color: Orange 5-TAMRA dUTP | Chr2:216224988-216402549 |
| CLN-1004 | Clone Library: RPCI-11<br>Clone Name: 954P20<br>Dye Color: Green 5-Fluorescein dUTP | Chr7:100405127-100578561 |
| CHR02-10-RE | Chromosome 02 Control Probe<br>Dye Color: Red 5-ROX dUTP | Chr2:93900000-96000000 |
